# Spatial and morphological organization of mitochondria across a connectome

**DOI:** 10.1101/2024.08.21.608912

**Authors:** Garrett Sager, Fabian Pallasdies, Robert Gowers, Snusha Ravikumar, Elizabeth Wu, Daniel Colόn-Ramos, Susanne Schreiber, Damon A. Clark

## Abstract

Neuronal function depends critically on the cell biological organization of mitochondria, which regulate calcium signals and produce energy, among other roles. However, little is known about how mitochondria are organized within the circuits of neurons that make up each brain. To uncover the systematic rules that govern mitochondria shape and position in a connectome, we analyzed the morphological and spatial organization of more than 100,000 mitochondria in over 1,000 visual projection neurons in the *Drosophila* connectome. We found that mitochondrial shape and size differ systematically between cell types, and are distinct enough between cell types to serve as an identifying fingerprint. Moreover, we derived three quantitative rules that describe how mitochondria are positioned within neurons relative to synapses and other subcellular features: (1) they are positioned with a precision of 2-3 microns; (2) their relative preference for pre- and postsynaptic sites and other subcellular features differs between axons and dendrites; (3) their positions were specialized to different cell types. These organizing rules correlated with functional and anatomical properties of the cells, including visual responses and input connectivity. We also find that, in the fly’s olfactory associative learning circuits, mitochondria are enriched at presynapses to particular postsynaptic cells by accumulating in functional sub-compartments of axons. Overall, our findings reveal a robust set of organizing principles for mitochondria within and between cells, uncovering cell biology that maps onto the organization of the connectome and adding new dimensions for understanding circuit function in the connectome.

## Main

Anatomical connectomes are reconstructed via nanoscale analysis of electron microscopy (EM) datasets that contain the cell biological features underlying neural circuit architecture. Inspections of these features have mainly focused on defining neuronal relationships at the level of the chemical synapse ^1,2^, but EM images also capture the relative positions, morphologies, and characteristics of the subcellular organelles that underpin synaptic function, including mitochondria. Mitochondria are organelles associated with critical physiological processes tightly linked to neuronal function, including but not limited to energy metabolism and calcium signaling in neurons ^3–7^. They are actively trafficked to synapses ^8–12^, and dysregulation in their localization and function disrupts synaptic function ^8,13–19^ and is associated with neurological disease ^20,21^. While mitochondria localization near synapses is well documented ^12,22–24^, we know little about their systematic distribution within and across neuron types, nor about how they associate with synaptic and other nanoscale structures within neuronal circuits. Understanding this organization within circuits would help reveal how cell biological features of neurons beyond the synapse—such as mitochondrial morphology and position—supports the function of neural circuits in the brain.

To investigate relationships between mitochondria and the organization of neural circuits requires data that (1) bridges scales from nanometers to 100s of microns, enabling understanding of the cell biological feature (the mitochondria) in the context of the circuit (connectome); (2) includes varying cell types of known function; (3) contains features (morphology, position, relationship, etc.) that can be extracted with sufficient representation for statistical power. EM connectomic data meet these requirements. The *Drosophila* hemibrain connectomic dataset ^1^ catalogs ∼25,000 neurons and their connections across roughly half of an adult fly brain. It is particularly well-suited to investigate the organization of mitochondria within neural circuitry, since entire circuits are represented, the mitochondria have been segmented ^25^, the specific cell types identified, and the chemical synapses annotated ^26^. This dataset includes 100s of times more neurons and mitochondria than were in most prior studies of mitochondrial morphology and placement ^27–29^.

To investigate the morphological and spatial organization of mitochondria in the context of circuits, we focused on lobula columnar (LC) visual projection neurons in the fly’s visual system (**Fig. 1a, b, S1a-e**) for three reasons. First, visual projection neurons have similar morphology, with well-defined dendrites receiving input in the lobula and axons making synapses in the ventrolateral central brain ^30–32^, connected via a neurite with few synapses, facilitating comparisons between neuron types and relationships between specific synapses and mitochondria. Second, specific cell types have between 20 to 100 cells in the circuit, providing statistical power to make comparisons within and between cell types. Third, the functions of these cells have been investigated, establishing their responses to specific visual features and their activation of specific behavioral responses ^30,31,33–40^, offering the potential to connect mitochondrial organization with known aspects of LC neural function.

**Figure 1.**
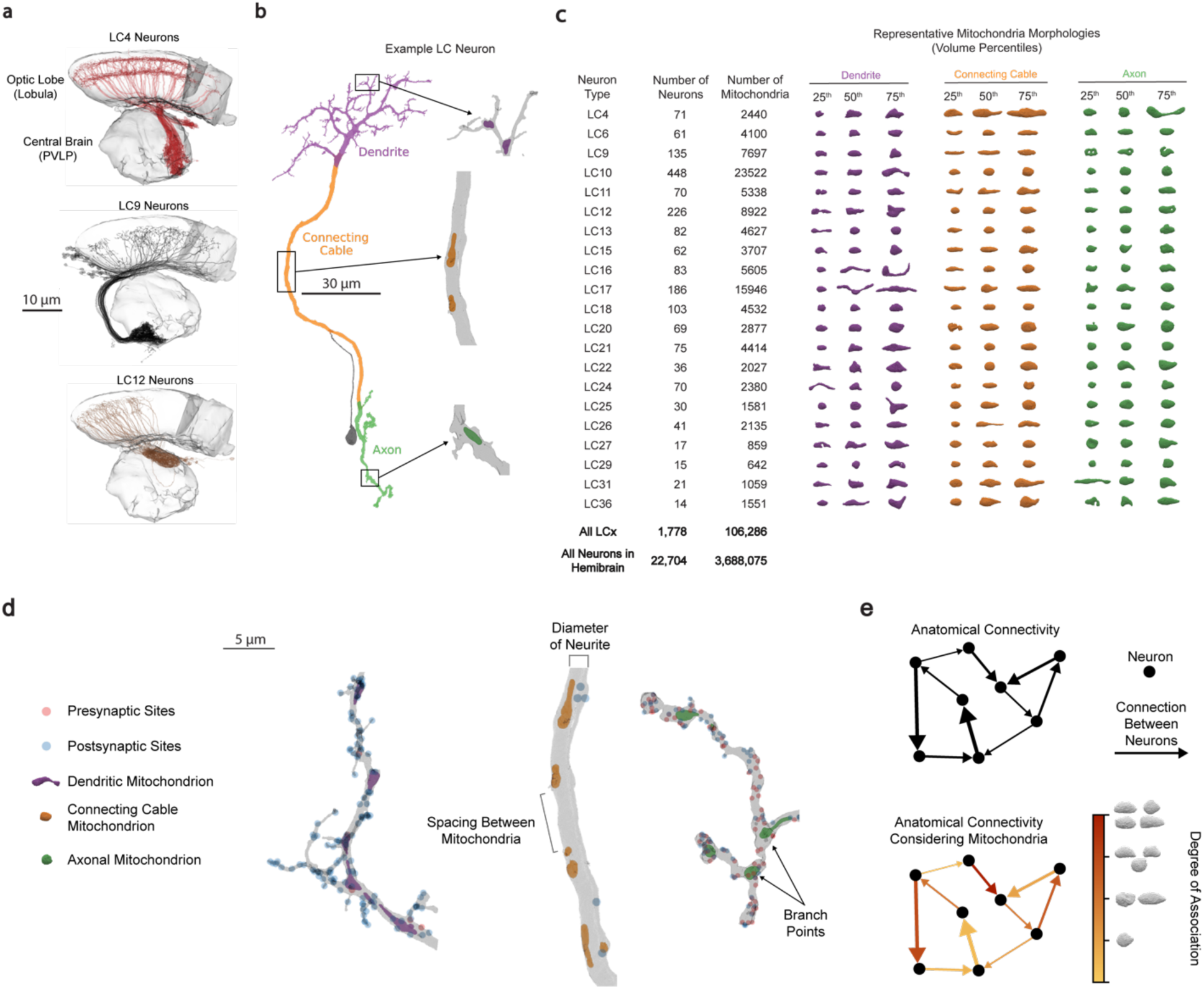
Visualizing mitochondria in LC visual projection neurons in *Drosophila*. (a) Renderings of LC4 (top), LC9 (middle), and LC12 (bottom) neurons with the lobula and PVLP neuropils. (Supplemental Table 1 contains URLs recreating neural renderings in many figure panels.) (b) Example LC neuron (bodyId 1218901359, neuron type LC4) with the dendritic arbor colored purple, the axonal arbor colored green, and the connecting cable colored orange. An enlarged section of each compartment is shown to visualize the mitochondria inside the neurite. (c) Representative mitochondria at the 25^th^, 50^th^, and 75^th^ volume percentiles are shown for each LC neuron type’s segments. To the left are listed he total number of neurons and total number of mitochondria across all neurons for each LC neuron type, all LC neurons types, and all identified neurons in the hemibrain. (d) Visualization of the LC4 neuron in panel (b) with points marking the locations of presynapses (red) and postsynapses (blue). Example features in the neurite (spacing between mitochondria, branch points, neurite diameter) are labeled. (e) Cartoon of an anatomical connectome between hypothetical neurons, with synapse strength represented by arrow width (top). Cartoon showing how network connections could be labeled by association with mitochondria (bottom).

In the hemibrain dataset, there are over 1700 segmented LC neurons containing over 100,000 segmented mitochondria of varying shapes and sizes in dendrites, axons, and connecting cables (**Fig. 1b,c**). Each mitochondrion is embedded within a neurite, establishing spatial relationships to presynaptic sites, postsynaptic sites, other mitochondria, and other features of the neurite (**Fig. 1d**). Since the neurons are components of identified circuits, mitochondrial locations are also associated with connectivity in the hemibrain connectome (**Fig. 1e**). In this study, we use these data to characterize mitochondrial shape within and between neurons, mitochondrial position within neurites and its relationship to function, and mitochondrial associations with specific synapses within circuits. The size and the high-dimensionality of this dataset required us to develop quantitative and statistical approaches to uncover the organization of mitochondria within circuits (see **Methods**).

### Mitochondrial morphology

To investigate how mitochondria are organized within the connectome, we first wanted to examine differences in mitochondria gross morphology across specific cell types. Mitochondrial shapes are diverse and likely affect key functional properties ^41^. Prior studies have shown that mitochondrial morphology differs between the axonal and dendritic segments (analyzing a few thousand mitochondria in a few dozen neurons ^29,42^), but it is not known how gross mitochondrial morphology varies across different neuronal cell types within an animal. To understand gross morphological differences, we plotted volume distributions and found they were different between neuron types and segments (**Fig. S1a,c**). While LC neurons have a range of volume distributions, some of the largest mitochondria exist in LC4 neurons. LC neurons encode many visual features ^30,34,39^, but LC4 plays a prominent role in detecting looming objects and initiating escape behaviors ^37,40^, so that larger mitochondria might reflect an energetic investment in a critical functionality.

Understanding the relationships between cell types and single-feature descriptors of mitochondria morphologies can be done by examining simple distributions. But to understand the relational features of the mitochondria, and to extract relationships that may not be apparent from visual inspection, it is necessary to embed the mitochondria into a multidimensional morphometric space, with each dimension related to a specific aspect of mitochondrial morphology: volume, aspect ratio, sphericity, etc. (**Fig. 2a**, **Fig. S2**, see **Methods**). To achieved this, we first embedded the LC mitochondria in a 21-dimensional morphometric space, then visualized this space using UMAP ^43^. This dimensionality reduction technique allows us to embed all mitochondria in two dimensions and color them by cell segment: dendrite, connecting cable, and axon (**Fig. 2b**). Our analyses revealed that the mitochondria in these different segments have different densities in this morphometric space, with longer, less round mitochondria more likely to be present in dendrites than in axons or connecting cables, consistent with prior reports ^29,42^ (**Fig. S3a**). To validate these differences, we trained a machine learning classifier to identify the segment type based on mitochondrial morphology (**Fig. 2c**). We prevented overfitting by 5-fold cross-validation and regularization (see **Methods**). The classifier correctly identified dendrites and axons 95% of the time, and connecting cables 80% of the time. Thus, systematic quantitative differences in mitochondrial morphology are strong enough to reliably classify neuronal segments such as dendrites, axons and connecting cables.

**Figure 2.**
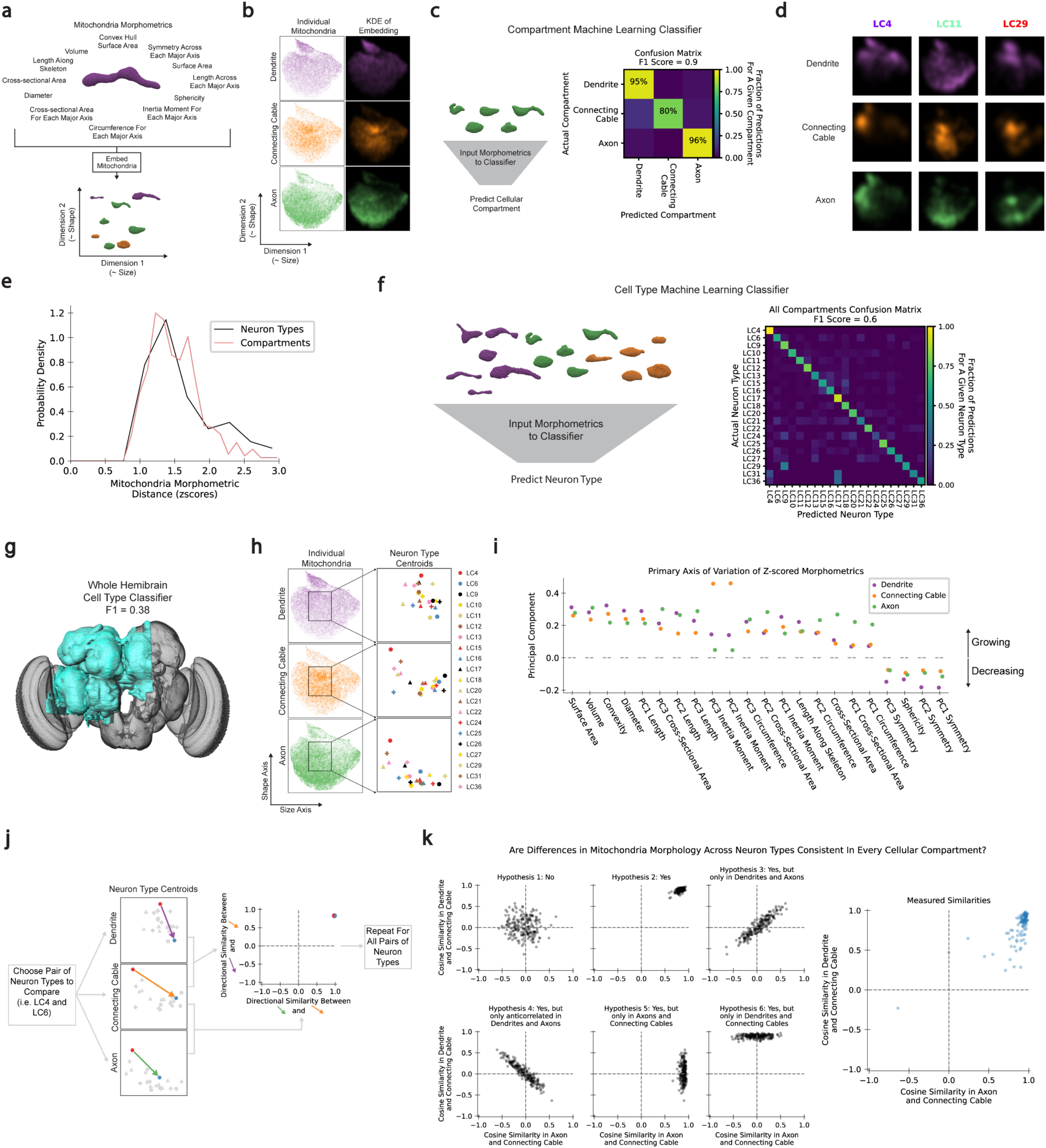
Mitochondrial morphology is specific to neuronal segments and neuron types. (a) Cartoon showing the 21 morphometric features used to embed mitochondria into a 2D plane using UMAP. (b) UMAP embedding of all mitochondria by their z-scored morphometric coordinates. In the plane, a kernel density estimate is calculated for each segment and colored by the respective segment’s legend color. (c) Cartoon of the random forest model predicting the cellular segment where a group of mitochondria are located. A confusion matrix for the predictions shows the true positive rate for each segment along the diagonal. (d) Kernel density estimates of the UMAP embedding for different segments of three example neuron types. Mitochondria map to distinct regions of the embedding when compared across compartments within a neuron or across neurons for the same compartment. (e) The probability density of dissimilarity in mitochondrial morphology across neuron types (peach) and across segments (black). The comparable distributions indicate that differences are as large between cell types as between cell segment types. (f) Cartoon of the random forest classifier that predicts the cell type based on the morphometrics of all the mitochondria in the dendrite, connecting cable, and axon (left). A Confusion matrix for the predictions is shown to the right. (g) When applied across the hemibrain, a random forest cell type classifier had an F1 score of 0.38 (see **Methods, Fig. S4**). (h) UMAP embedding of all mitochondria by their z-scored morphometrics, as in panel (b), but highlighting the centroid location of the mitochondria in different neuron types. (i) First principle component loading of the morphometrics for all mitochondria in the dendrites (purple), connecting cable (orange), and axons (green), showing the dimension of most variance in the mitochondrial morphology. (j) Cartoon illustrating how differences in mitochondria morphology between cell types were compared across cellular segments (see **Methods**). The directional similarity was computed by using the cosine similarity. (k) Cosine similarities of the difference in centroid locations in the full, 21 dimensional morphometric space between all pairs of neuron types. Various hypothetical outcomes are shown in the first five plots illustrating the outcome if the cell type specificity of mitochondria morphology were (hypothesis 1) not similar across neuron segments, (hypothesis 2) similar across all neuron segments, (hypothesis 3) only similar in axons and dendrites, (hypothesis 4) only anti correlated in axons and dendrites, (hypothesis 5) only similar in axons and connecting cables, or (hypothesis 6) only similar in dendrites and connecting cables. The measured similarities are shown in the far right.

We next examined differences in mitochondrial morphology between neurons (**Fig. 2d, 2e**). Strikingly, we found substantial differences across neurons for each segment (**Fig. 2d**). For instance, LC4 and LC11 have notably distinct mitochondrial morphology in their dendrites and axons, with LC4 mitochondria tending to be larger and less spherical. Such differences exist between many pairs of neuron types, in axons, dendrites, and connecting cables (**Fig. S3b**). To evaluate the size of this neuron-to-neuron variability, we computed a distance metric within neurons between segments and compared it to the same metric within segments between neurons (**Fig. 2e**). Interestingly, the differences between neuron types are as large or larger than the differences between segments. This suggests that neuron specific rules govern mitochondria morphology within neurons.

Based on these differences between neurons, we hypothesized that cell types might be identifiable merely by the morphology of their mitochondria. To test this, we again trained a simple classifier, but this time to predict neuron type (see **Methods**). Again, we prevented over-fitting by 5-fold cross-validation. When the mitochondrial morphology from dendrites, axons, and connecting cables were used, the classifier identified neuron types with an accuracy F1 score of 0.6, compared to an F1 score of 0.05 for a classifier guessing at random (**Fig. 2f**). The classifier could also identify neurons types with substantial accuracy based on dendritic, connecting cable, or axonal mitochondrial morphologies individually (**Fig. S3c**). Training this classifier across all neuron types in different neuropils in the dataset, beyond just LC neurons, yielded an accuracy F1 score of 0.38, compared to 0.04 for random guessing (**Fig. 2g, Fig S4**, see **Methods**).

When neurons have distinctive mitochondrial morphologies, what is the axis along which they vary? To answer this question, we first found the average mitochondrial morphology for each cell type and segment (**Fig. 2h**). We then computed how these morphometrics varied across cell types (**Fig. 2i, Fig. S3d**). The first principal component accounted for ∼50% of cell type-to-cell type variability across each of the cell segments, and was similar for dendrites, axons, and connecting cables. It corresponded to mitochondria increasing in volume while becoming less spherical and more complex in shape (**Fig. 2i**). We next asked whether differences between cell types were coordinated across segments, or whether different segments could vary independently. We therefore computed the difference in mean mitochondrial morphology between cell types within each segment (**Fig. 2j**). We found that the differences between segments of a given pair of neuron types tended to be in the same direction, so that the shift in morphometric space was in the same direction for dendrites, axons, and connecting cables (**Fig. 2k, Fig. S3e**). Our data is consistent with correlated changes in mitochondria morphology across segments, potentially reflecting cellular-level control of mitochondrial morphology across segments.

Overall, this analysis demonstrates (1) that mitochondrial shape differs systematically between neurons, with as much variation as between segments; (2) that mitochondrial morphology might be a productive means of identifying neuron types within truncated neurons in EM databases; and (3) that there are neuron-specific rules that define features of mitochondria morphology that are coordinated across their dendrites, axons, and connecting cables.

### Mitochondrial positioning

Are there rules governing mitochondrial position within neurons? To address this question, we fit a model to predict how mitochondrial position is affected by neurite properties and features along the neurite length (**Fig. 3a-b**). Prior studies have suggested that mitochondria collocate with presynaptic sites ^12,20,22,25^, with postsynaptic sites ^44,45^, and with neurite branch points ^46,47^. Here, we fit a quantitative model that describes how specific features, including distance to synapses and other mitochondria, neurite diameter, and number of supported terminal branches, among others, influence mitochondria positioning (**Fig. 3c**). We chose a logistic model to predict the likelihood of a mitochondrion at a given location, relative to a control of shuffled mitochondria within the same cell and segment (**Fig. 3c**, see **Methods**). Logistic models have two advantages here: (1) the model parameters can be easily interpreted in terms of log likelihoods and (2) they can account for correlations among the many simultaneously examined features, which builds on prior experiments that tended to examine one feature at a time. In particular, by accounting for the correlations between features, models are prevented from attributing likelihood to spurious features. By making comparisons to a shuffled control, we ensure that the model identifies positioning rules, rather than just differences in features between cells and segments. We prevented over-fitting by five-fold cross-validation and lasso regression.

**Figure 3.**
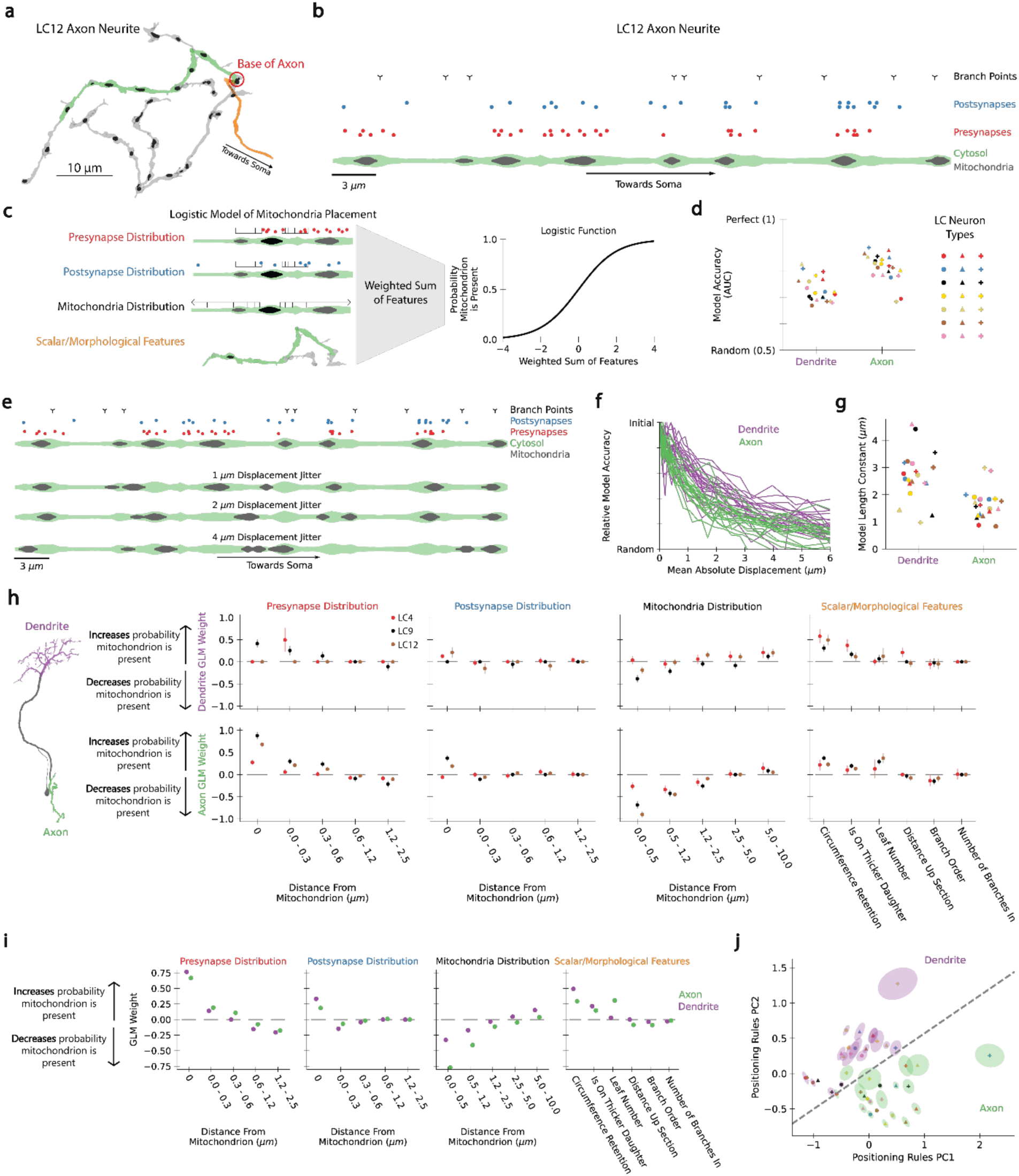
Quantitative positioning rules show the precision and pattern of mitochondrial localization. (a) An example LC12 axon (bodyId 1750630587) showing mitochondrial positions. The base of the axon is circled in red. The green portion of the neuron is used in the cartoon in panel (b). (b) An example section of the axon from panel (a) with the cytosol colored green and the mitochondria colored grey. The locations of the presynapses (red circles), postsynapses (blue circles), and branch points (black branch symbol) are indicated. The soma is to the right. (c) Cartoon of the logistic model for mitochondrial positioning. Input features describe the presynapse spatial distribution, postsynapse spatial distribution, neighboring mitochondria spatial distribution, as well as various scalar and local neuronal morphology features (see **Methods**). A weighted sum of these features is then converted by a logistic function to the probability that a mitochondrion is present. The weights are optimized to predict true mitochondrial positions relative to randomly placed mitochondria (see **Methods**). (d) The model accuracy (AUC) of the logistic positioning model for each neuron type’s dendritic and axonal arbor. Neuron types are labeled as in Fig. 2h. (e) Cartoon of the same axonal segment from panel b after the mitochondria have been jittered with mean absolute displacement of 1 *μm* (top middle), 2 *μm* (bottom middle), or 4 *μm* (bottom middle). (f) The logistic positioning model accuracy decreases as the mitochondria are jittered up to 6 *μm*. All LC neuron types are shown for the axonal and dendritic logistic positioning models. The initial model accuracy is scaled to match to permit comparisons between different neurons and segments, while the random model accuracy corresponds to an AUC of 0.5. (g) The length constant in microns for the functional forms in (f). Each curve was fit to an exponential decay, *e*^−*x*/*λ*^, where *λ* is the length constant. Neuron types are labeled as in Fig. 2h. (h) Plot of logistic model weights for three example neuron types. Error bars show the 95% confidence interval. The features are defined in the **Methods**. (i) Plot of logistic model weights for dendrites and axons, combining data from all neurons. Error bars show 95% confidence intervals, but are smaller than the points. (j) Logistic model weights projected into their first two principal components. Ellipses represent one standard deviation in parameter confidence in the plane. Ellipses are colored by dendrite vs. axon; each point’s color and shape represent neuron type, as in Fig. 2h.

We trained different models on dendritic and axonal mitochondria in all 21 LC neuron types in our study. The trained models could predict the positions of true mitochondrial locations compared to shuffled ones in both axons and dendrites across many cell types (**Fig. 3d**, see **Methods**). We note that models succeeded even though mitochondria can be both mobile and docked ^8–11,13^. We hypothesize that the accuracy of these models is dominated by mitochondria that are predictably docked at specific positions relative to the features we examined, suggesting that a substantial fraction of the mitochondria are docked.

To quantify the precision of mitochondrial positioning within dendrites and axons of these cells, we randomly displaced (or jittered) the positions of all mitochondria in our dataset by different distances ranging from 0 to 6 um (**Fig. 3e**). We then asked how well the original models predicted the displaced mitochondria. If the model had learned a very precise positioning rule, then it would not be able to predict mitochondrial placement when even small displacements were added. On the other hand, if the model had learned a broader probability distribution, then the small added displacements would not much affect its accuracy. We plotted the relative model accuracy as a function of mitochondrial displacement (**Fig. 3f**) and found that these accuracy curves decayed to random prediction with displacements over a few microns. We quantified the length scale over which this decay occurs by fitting the curves to exponential functions (**Fig. 3g**). This analysis demonstrates that mitochondria are positioned with 2-3 um precision in both axons and dendrites, with slightly higher precision in axons.

Next, we examined the models themselves to understand the rules by which mitochondria are positioned in dendrites and axons of each cell type. We investigated how different features affect mitochondrial likelihoods by examining the weight put on features at different distances away from each location (**Fig. 3h, Fig. S5c)**. In dendrites and axons, presynapses tended to increase the probability of a mitochondrion, in agreement with prior studies ^12,22^. Directly adjacent synapses increase the probability of finding a mitochondrion (“0 distance” weights are positive), but nearby synapses can also increase the probability of finding mitochondria, to a range of ∼0.5 microns. Postsynapses increased mitochondrial probabilities to a smaller degree, while other mitochondria within a couple microns decreased the probability of finding a mitochondrion, especially in axons. Among the scalar features, larger neurites and being in the larger of two daughter branches both increased the probability of a mitochondrion in dendrites as well as axons (**Fig. 3h**), consistent with neurite morphology influencing mitochondrial transport ^48^. The positioning rules are quantitative representations of the data, showing not just correlations with single features but picking out the most important features that predict mitochondrial position. While prior work had shown that mitochondria were associated with presynapses, postsynapses, and developing branches ^8–12,46,47,49^, these fitted models quantitatively describe those relationships and disentangle the contributions of individual features by decorrelating the different features from one another.

While there were broad trends in the positioning rules, they differed across both neuron type and segment type (**Fig. 3i,j, Fig. S5d,e**). The dendritic and axonal positioning rules were most distinct in (1) how axonal mitochondria ‘repelled’ one another more strongly than dendritic ones and (2) by the influence of the number of leaves (terminal branches) on axonal mitochondrial positioning (**Fig. 3i**). Differences between positioning rules were visualized by projecting the fitted model for each neuron segment onto the first and second principal components of the set of all model parameters (**Fig. 3j**). The dendritic and axonal positioning rules are clearly distinct in this visualization, clustering below and above the dashed line. This effect cannot be accounted for solely by segment-specific differences in pre- and postsynapses, since both axons and dendrites contained both pre and postsynaptic sites. Moreover, this visualization also clarifies that different neurons have specialized positioning rules.

### Correlations with neural function

Our next goal was to understand whether mitochondrial positioning correlated with functional and connectivity properties in the brain. We began by asking whether mitochondrial density and positioning in different regions of the brain correlated with spontaneous activity that had been measured in prior studies ^50,51^. To do this, we mapped the number density of presynapses and number density of mitochondria over brain regions in the hemibrain (**Fig. 4a**, **Fig. S10a,b,c**, see **Methods**). These densities were characteristic of each brain region, so that they were correlated across brain hemispheres (**Fig. 4b**). We next correlated these structural properties of brain regions with spontaneous calcium, ATP, and pyruvate signals recorded in the different regions ^50,51^. Since the mean indicator signal depends on reporter expression strength, which varies between brain regions, we quantified the activity as the standard deviation of calcium signals, which tend to reflect deviations from baseline activity (**Fig. 4c, Fig. S6a, Fig. S9b**). We then plotted this response metric for each indicator against the presynaptic density, mitochondrial density, mitochondrial volume per synapse, and fraction of presynapses associated with mitochondria for all central brain neuropils (**Fig. 4d**). The calcium activity and ATP activity were significantly correlated with the fraction of presynapses associated with mitochondria, while pyruvate indicator showed minimal correlations (**Fig. 4d, Fig. S6b, Fig. S9b**). This correlation could arise because mitochondria enhance synaptic strength ^18^, generating more strongly reinforcing activity, or because synaptic activity attracts mitochondria ^12,49,52–55^, so that the mitochondria could reflect more synaptic activity. Importantly, our analyses uncover a correlation between the association of mitochondria and presynaptic sites, and mitochondria and known ATP states in neurons.

**Figure 4.**
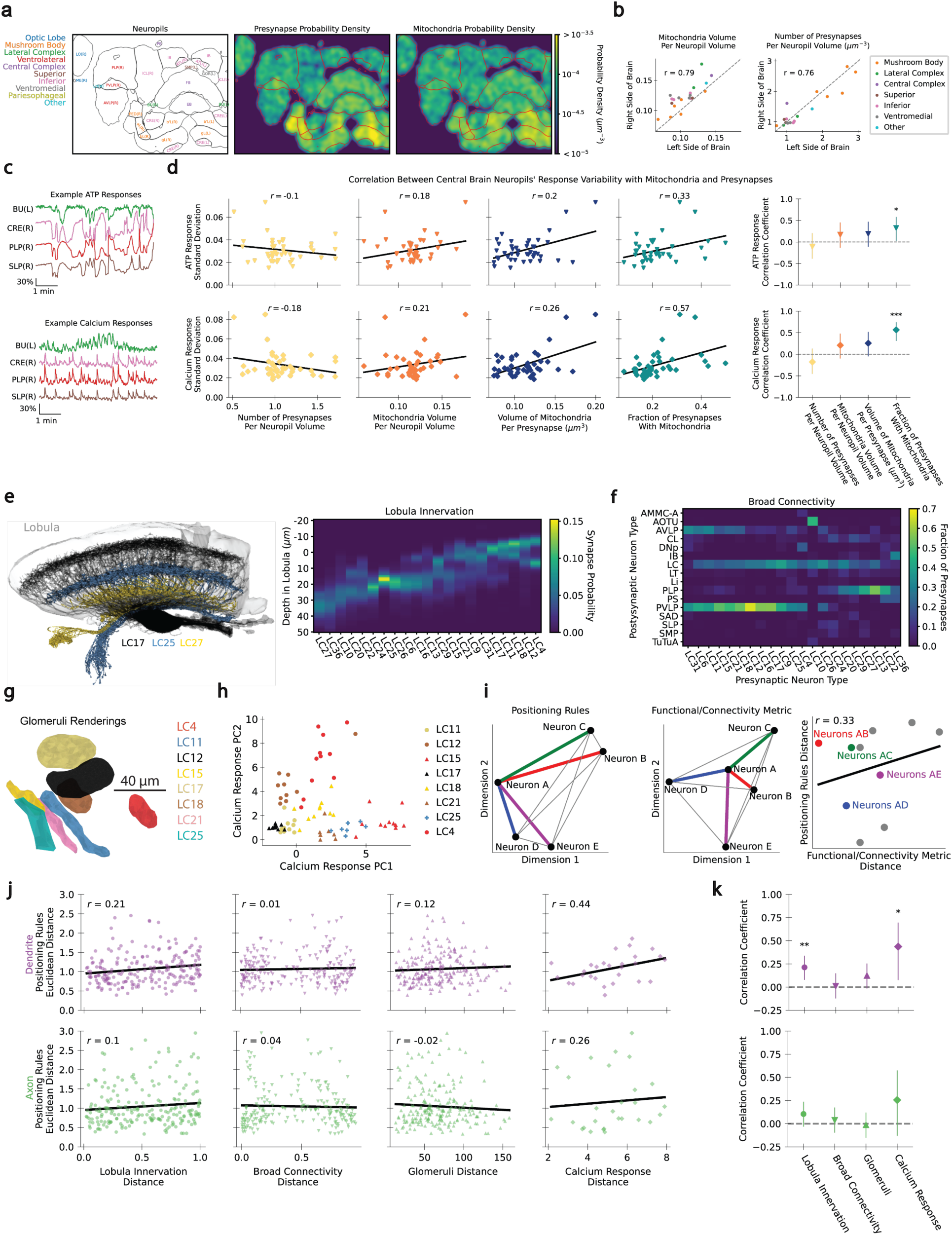
Mitochondrial positioning correlates with spontaneous activity and with visually evoked activity. (a) Cross-section of the fly brain with labeled neuropils (left). The density of presynapses (middle) and mitochondria (right) in the slice are calculated using kernel density estimation. (b) Scatter plots comparing mitochondrial and presynaptic densities in neuropils in the left and right halves of the brain, where both exist in the dataset. The unity line is shown in gray. (c) Example ATP and calcium indicator traces for four central brain neuropils (data from ^50,51^). (d) Scatter plots of the standard deviation in the ATP and calcium indicator responses for all recorded central brain neuropils against the number of presynapses per neuropil volume, total volume of mitochondria per volume of neuropil, total volume of mitochondria per presynapse, and fraction of presynapses with mitochondria. Best fit lines are shown in black. The Spearman correlation coefficients for each plot are computed with 95% confidence intervals (right). (e) Visualization of three neuron types, highlighting their projections in the lobula (left). Kernel density estimates of every LC neuron’s innervation pattern along the depth of the lobula (right). A link recreating the rendering is in Supplemental Table 1. (f) Adjacency matrix of the fraction of connections from each presynaptic LC neuron type to each postsynaptic neuron type. (g) Visualization of the glomeruli (axonal synapses) for each LC neuron type analyzed in ^39^. (h) First two principal components of calcium responses to many different visual stimuli. Data from ^39^. (i) Cartoon for how positioning rule distance may be correlated with distances in function or connectivity of the neuron. (j) Scatter plot of positioning rule Euclidean distance with each functional and connectivity distance for all pairs of LC neuron types. Black line represents the best fit line. (k) Spearman correlation coefficients from panel (j) with 95% confidence intervals. P-value significance is represented by asterisks (* < 0.05, ** < 0.01)

Could mitochondrial positioning rules in different neuron types be related to functional differences between neuron types? To address this question, we compared the mitochondrial positioning rules measured in identified LC neurons (**Fig. 3**) to the connectivity and measured functional properties of the LC neurons. We first collected a variety of features that relate the functional properties of LC cells: First, LC cells innervate different layers within the lobula, where they receive inputs from functionally distinct inputs ^56,57^ (**Fig. 4e**). Thus, the synaptic probability at different depths in the lobula represents the distribution of neuron types that provide input to each LC. Second, looking at downstream connectivity, different LC neurons synapse onto different broad classes of postsynaptic partner types (**Fig. 4f**). Third, glomerular positions create a map in which physical proximity is related to functional properties ^39^ (**Fig. 4g**). Last, responses of different LC neurons to a suite of visual stimuli have been systematically measured ^39^, providing an abstract space for measuring the functional similarity between different neuron types (**Fig. 4h**).

To assess how mitochondrial positioning rules relate to these different connectivity and functional features of the LC neurons, we next computed a distance between the positioning rules for each pair of neurons, based on the distance between their fitted parameters in the logistic regression (**Fig. S6a-d**). The neurons may be placed in both a positioning rule space and in the space of other connectivity and functional features (**Fig. 4i**). If the spaces are related to one another, similar neurons will be clustered in both spaces, and dissimilar neurons will be distant in both spaces. We can measure the degree of similarity by computing distances between all pairs of neurons in the positioning space and correlating them with distances between the same pairs of neurons, computed from connectivity and functional metrics (**Fig. 4i**).

We therefore plotted the positioning rule distance against the lobula innervation distance, the broad postsynaptic connectivity distance, the glomerular distance, and the calcium response distance (**Fig. 4j, S6e**). Interestingly, we found significant correlations between the dendritic rules and the dendritic arborization, and between the dendritic rules and calcium responses (**Fig. 4k**), and even stronger correlations exist when assessing rule predictivity rather than parameter differences (**Fig. S6e**). These correlations mean that neurons with similar positioning rules in their dendrites also share similar laminar innervation in the lobula and similar calcium responses to visual stimuli. While the causal mechanism underlying this correlation is not known, this correspondence suggests that local mitochondrial placement is linked to and could influence neuronal function, potentially mediated by known mitochondrial roles in calcium signaling and energy requirements at synapses ^44,58–61^. An alternative, non-exclusive explanation is that the dendritic morphology and innervation of LC neurons strongly influences both positioning rules and their functional properties, since mitochondria positioning depends on synaptic activity^12,49,52,54,55^.

### Dependence on postsynaptic partners

In the analysis so far, we have treated individual presynapses as anonymous, that is, without considering the identity of their postsynaptic partner. We wondered whether mitochondria are preferentially associated with specific presynapses (as determined by their postsynaptic partner identity), indicating, for example, functional differences across specific synapses and their association with mitochondria. To answer this question, we first defined a synapse as being associated with a mitochondrion if the synapse is adjacent to any part of the longitudinal extent of that mitochondrion (**Fig. 5a**, see **Methods**). We will refer to the fraction of synapses associated with a mitochondrion as ‘mitochondrial coverage’. We then examined the degree of enrichment of mitochondrial coverage for each neural connection by measuring whether specific presynapses were more or less likely than average to be with mitochondria, conditioned on the postsynaptic target (**Fig. 5b**). In LC4 neurons, presynapses to some postsynaptic cells were enriched and some depleted, but the effect sizes were relatively small and the effects not significant (**Fig. 5c**); similar results held for other LC neurons (**Fig. S7**).

**Figure 5.**
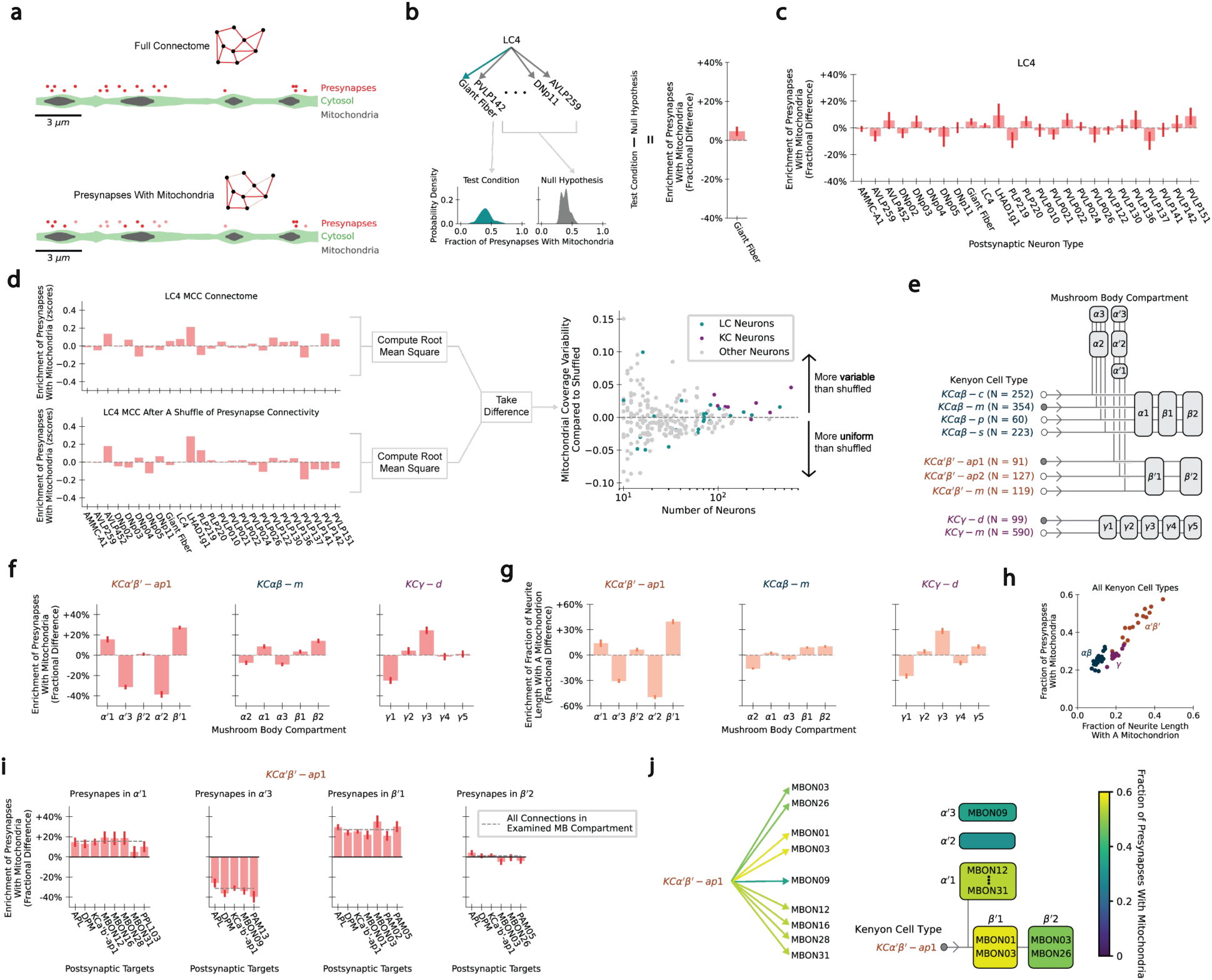
Cells in an associative learning center enrich mitochondria at the presynapses of specific output neurons. (a) Cartoon of the full connectome (top), repeated showing only presynapses associated with mitochondria (bottom). Presynapses not associated with mitochondria are set to transparent. The fraction of presynapses that are associated with mitochondria is the presynaptic mitochondrial coverage, which can also be specified for subsets of presynapses, for instance presynapses to a particular postsynaptic neuron. The neurite segment is the same LC12 axonal neurite as in Figure 3. (b) Demonstration of how to compute the enrichment in mitochondrial coverage of LC4 presynapses onto the giant fiber. The enrichment for one postsynaptic neuron type is computed compared to all other presynapses. (c) Bar plots show the enrichment in presynaptic mitochondrial coverage for each postsynaptic neuron type. (d) Cartoon showing how we compare the distribution of presynaptic mitochondrial coverage to a null hypothesis of shuffled synaptic connectivity. The difference in the root mean square mitochondrial coverage before and after shuffling synaptic connectivity defines the deviation of the measured coverage from random. Positive and negative values indicate more variable and more uniform coverage than random. Right: A scatter plot of the relative variability against the total number of neurons in each neuron type. We considered only neuron types with at least ten neurons. (e) Cartoon of the Kenyon cell (KC) types innervating mushroom body compartments. The filled in circles represent the three KC types used as examples in later panels. Arrows indicate the direction of information flow (dendrite to axon). (f) Enrichment of presynapses with mitochondria in each functional compartment. Enrichment was computed as in (b), but the test condition was all presynapses in the specified mushroom body compartment, while the null hypothesis was all the other presynapses. (g) Enrichment in neurite length with a mitochondrion present (i.e., linear mitochondrial density, see **Methods**) in different compartments. This is computed as in (f) but using linear mitochondrial density rather than presynaptic mitochondrial coverage. (h) Presynaptic mitochondrial coverage plotted against linear mitochondrial density for each mushroom body compartment and each KC type. KC classes are as labeled. (i) Bar plots showing the enrichment of presynaptic mitochondrial coverage for neuron types postsynaptic to *KCα*′*β*′ − *ap*1. Enrichment is for the identified presynapses compared to presynapses in the other mushroom body compartments. The grey dashed lines represent the enrichment in each compartment, as in panel (f). (j) Left: connectivity diagram using the color of arrows to represent *KCα*′*β*′ − *ap*1 presynaptic coverage of different mushroom body output neurons (MBONs). Right: cartoon of presynaptic coverage by mushroom body compartment, with MBONs labeled. The color legend is the same in both cartoons.

We therefore examined the entire hemibrain to see whether other neuron types (for which there exist more than 10 neurons in the hemibrain) showed stronger enrichment. To do this, we first computed the variance in coverage across presynapses with different target neurons, which we compared to a null hypothesis in which synapse identity was shuffled (**Fig. 5d**). We found that Kenyon cells (KCs) had consistently larger variability than the null hypothesis (**Fig. 5d**). The large number of each KC type made the differences in coverage statistically highly significant (**Fig. S10d**). Mitochondria are critical for synaptic plasticity ^44,45,62^, and the KCs in the mushroom body are a major location for plasticity and memory formation in the fly brain (**Fig. 5e**). Nine populations of Kenyon cells (KCs) contain cells that are each activated by different specific odors and make synapses onto mushroom body output neurons (MBONs), which guide aversive and appetitive behaviors ^63–65^. The plasticity of these KC-MBON synapses is a site for associative learning. We therefore decided to ask whether mitochondria positions in KCs are related to presynapses with specific postsynaptic targets associated with these known associative learning and plasticity functions in KCs.

Next, we focused on the subcellular localization of mitochondria within KCs in the context of the connectome. The mushroom body is divided into 15 compartments that form functional units (**Fig. 5e**), so that each KC axon passes through several compartments, making contacts with MBONs that each enervate a single compartment ^64^. We therefore examined the distribution of mitochondrial coverage of presynapses in the different functional compartments over the different KC types (**Fig. 5f, S8a,h**). Within a single KC type, mitochondrial coverage varied substantially between compartments, with enrichment often exceeding half the standard deviation (**Fig. S8a**). Presynapses in some compartments could be associated with mitochondria up to 30% more or 40% less than in other compartments, in the same KC type. Thus, single KC types have different mitochondrial coverage statistics within different compartments (and hence postsynaptic partners), which could relate to the differential roles played by different KCs in learning ^66^.

We wanted to know if this coverage difference was attributable to different densities of mitochondria in the different compartments. We therefore computed the linear density of mitochondria as the fraction of the neurite length with a mitochondrion present. We analyzed the distribution of this density within the different compartments of different KC types (**Fig. 5g, S8b**), finding that within a KC type, the compartments with more mitochondrial coverage also had higher mitochondrial densities. We plotted mitochondrial coverage against the density, revealing distinct clusters of KCs with different linear densities and mitochondrial coverage (**Fig. 5h**). The high density of mitochondria in the *α*′*β*′ KCs is consistent with their known higher firing rates ^67^ and odor responsiveness ^68^, revealing a reasonably linear relationship in different KC types (**Fig. 5h, S8d**). Importantly, the relationship differed quantitatively across cell types (**Fig. S8e**), indicating that different cells use different rules to map the linear density into synaptic coverage, potentially related to their functional differences.

Since the mitochondrial coverage of presynapses in KCs differed between compartments, we wondered if the coverage was different between postsynaptic cell types within compartments. We therefore plotted the coverage of presynapses within compartments, conditioned on the postsynaptic cell type (**Fig. 5i, Fig. S8f**). Across compartments and KC types, all cell types had presynaptic mitochondrial coverage at roughly the same rate within a compartment, but each compartment could be substantially different, as we had seen before (**Fig. 5f**). One cell named APL, a large inhibitory neuron that is not an MBON, receives inputs from all KCs in all compartments. For this neuron, we found that the presynaptic mitochondrial coverage varied by compartment, matching the overall compartmental coverage rule (**Fig. S8g**). These data thus support a model in which the probability that a mitochondrion is associated with a presynapse depends on the functional compartment it is in. This regulates mitochondrial coverage of presynapses to specific postsynaptic neurons because they can be compartment specific (**Fig. 5j**). It is not simply that there are more mitochondria in compartments closer to the soma; for instance, KC*γ*-d cells have lower synaptic coverage in compartment *γ*1 than in *γ*3, which is further from the soma. Mitochondria dynamics increase in mushroom body compartments during long term memory formation ^55^, and our results lead us to hypothesize that mitochondria could be dynamically enriched in different compartments during associative learning, placing them where their services are most needed.

## Discussion

Since the earliest studies by George Palade, electron microscopy has sought to map functional properties onto physical structures. New connectomic data has ushered in opportunities to discover new principles of cell biological organization and link them to circuit structure and function. As data and segmentation have improved, new questions relating subcellular structures and morphology to network connectivity and neuronal function can be addressed. Here we focused on mitochondria, quantitatively examining millions of these organelles and their relationship to other cell biological features to obtain precise rules describing their morphology and location in neurons. These analyses revealed three main principles: (1) Mitochondrial morphology and position are cell type specific (**Fig. 2, 3**). In fact, mitochondrial morphology can be sufficiently distinct between neuron types to serve as an identifying fingerprint that enables cell typing even when neuronal morphology is incomplete (**Fig. 2**), as it often is in EM datasets. (2) Mitochondrial position is related to functional properties of the neural network (**Fig. 4**). Relationships between synapses and mitochondria correlated both with spontaneous, regional activity as well as with visually evoked activity across single neuron types. (3) Mitochondria occupy positions related to specific synaptic connections in the network (**Fig. 5**). We found that mitochondria can be enriched at specific presynapses by increasing their presence in functional compartments with specific postsynaptic targets (**Fig. 5**), an organization that could represent an efficient general mechanism for regulating demands mediated by mitochondria.

The organizing principles uncovered here are likely to generalize to other circuits and to influence the functional properties of neural circuits. Within neurons, mitochondria are transported along microtubule networks to different subcellular locations, especially to synapses ^9–11^. There, they support synaptic integrity and affect synaptic strength and plasticity ^8,13–19^, so that their morphology and location impacts cellular and circuit function. Critically, however, the position of mitochondria is also regulated by neuronal and synaptic activity and associated factors like calcium and ATP concentrations ^12,49,52–54,69–75^, likely to meet the metabolic demands of synaptic transmission ^76^ and related processes like the formation of long-term memories ^55^. If mitochondrial location and morphology reflect on-going activity, then the anatomical snapshot of the nervous system captured in connectomic datasets contains an imprint of past neural activity. Such imprinting would provide one explanation for the observed correlations between mitochondrial positioning and neural functional properties (**Fig. 4**). To better understand potential imprints of neural activity, it will be important to investigate how cellular mechanisms interact locally to position and shape mitochondria within a dynamic circuit environment.

The broad roles for mitochondria in neural circuits mean that they can add a latent functional dimension to the connectome, adding to existing features of cell morphology, synapse location, and neurotransmitter type. As the varied functions for mitochondria at different synapses become better understood, their positioning and morphology within the connectome can provide complementary information that can be overlayed on top of the anatomical connectivity to better understand functional connectivity in the brain. Mitochondrial organization within cells and circuits could also be used to further categorize neurons, synapses, and functional compartments.

Our study relates the morphology and position of mitochondria to the structure of neurons in circuits. It adds to work analyzing other subcellular structures in large EM datasets ^77–80^. Large EM datasets make possible analyses and comparisons that are not possible with smaller datasets, but the richness and complexity of these datasets both invite and necessitate quantitative analyses. Some of the quantitative approaches developed here could be extended to other subcellular structures beyond mitochondria. Machine learning approaches, used elsewhere to identify neurotransmitter types ^81^ and segment volumetric data ^82^ and used here to classify cells and identify rules for localizing mitochondria, seem likely to prove particularly useful. As it becomes possible to analyze increasingly rich subcellular relationships—for instance, by including abundance and positions of varying vesicular populations, the presence of autophagosomes at the synapse, or endoplasmic-reticulum contact sites—quantitative structural approaches like the ones applied here could be used to discover new principles of cell biological organization and how cellular structure connects to circuit function.

## Acknowledgements

We thank S. Bahmanyar, D. Breslow, J. Gendron, A. Kuan, A. Mallya, J. Musser, L. Scheunemann for helpful comments and suggestions on this research. GS was supported by a Fulbright Fellowship and a NSF GRFP. Research in DAC-R’s lab was supported by NIH/NINDS grant R35 NS132156-01. Research in SS’s lab was supported by the Einstein Foundation Berlin grant EP-2021-621. This research in DAC’s lab was supported by NIH R01 EY026555.

## Contributions

GS, FP, RG, SS, and DAC conceived analytical and computational approaches. GS, FP, and RG performed the research. GS and EW visualized data. SR and DC-R visualized and interpreted data. GS, FP, SR, DC-R, SS, and DAC wrote and edited the paper.

## Methods

### Software

A yml file with the packages used is included in the GitHub repository ^83^. All code to recreate these analyses is available at: [Github link to come with publication].

### Choosing neuron types

For most of the study, we chose to investigate LC neurons because there are many types, each with approximately 100 neurons per type. All LC and Kenyon cell types with at least 10 individual instances are analyzed.

### Processing neuron skeletons of LC neurons

The default neuron skeletons contained inaccurate measurements of the radius of the neuron and many false positive detections of branches in thick parts of the neuron. We recomputed the radius measurements by iterating through all nodes in the skeleton and finding a plane that minimizes the cross-sectional area (**Fig. S10**). The radii in the dataset were up to 50% different than the newly computed radii (**Fig. S10**). The normal vector of the plane with minimal cross-section also defines the instantaneous orientation of the neuron. Other features were stored for later analyses: the area of mitochondrial tissue intersected by the cross-sectional plane, the circumference of the neuron, and the unit vector of the plane’s normal vector (in spherical coordinates).

To identify false branches in skeletons, 1028 branches were manually labeled as true or false branches. The branches were selected uniformly across LC neuron types to ensure equal representation of each LC neuron type. We trained a decision tree to predict true branches and discovered that classifying branches by whether they are at least 91.4 nm longer than their width resulted in an 84% true positive accuracy for identifying true branches. Consequently, we used this threshold and removed from the skeleton all branches that are not at least 91.4 nm longer than their diameter.

Since the hemibrain dataset crops the optic lobe, many LC neurons have a cropped dendritic arbor, missing cell body, and/or cropped axon (**Fig. 1a**). We examined each LC neuron and manually labeled whether the neuron contained an axon, dendrite, and/or soma (**Fig. 1b**). For neurons with a cell body, the root node of the skeleton was defined at this point by finding the node with the largest radius. For neurons missing a soma but containing both arbors, the root node was defined as a skeleton node that on one side only contains optic lobe synapses and on the other side only contains synapses in the central brain. For neurons missing a soma and only containing one arbor, the root node was the farthest skeleton node from the median position of the central brain or optic lobe synapses, depending on whether the neuron had an axon or dendrite, respectively.

Each node of the skeleton was then classified as being in the axon, dendrite, soma, connecting cable, or cell body fiber. The connecting cable is the long piece of the neuron connecting the axon and dendrite, while the cell body fiber is the long piece of neuron that connects the soma to the connecting cable. Each arbor is defined as the part of the neuron containing synapses in the optic lobe (dendrite) or central brain (axon). To define the dendrites, we started at the skeleton node closest to the median position of all optic lobe synapses and moved in the direction of the root node one node at a time until at least 98% of optic lobe synapses’ closest skeleton nodes were in the direction away from the skeleton root node. That node defined the end of the dendrite. To define the axons, we started at the skeleton node closest to the median position of all central brain synapses and moved in the direction of the root node one node at a time until at least 98% of central brain synapses’ closest skeleton nodes were in the direction away from the skeleton root node. That node defined the end of the axon. The connecting cable nodes were then defined as the set of nodes that connect the base node of the axon and dendrite. If the neuron does not contain an uncropped axon and dendrite, it will not have a connecting cable. Cell body fiber nodes were defined as the set of nodes connecting the soma to the connecting cable. The node connecting the connecting cable and cell body fiber is referred to as the main bifurcation node. Soma nodes were defined by starting at the node with the maximum radius, and moving one node at a time towards the main bifurcation node until the radius is less than the median radius of the initially defined cell body fiber.

### Saving features for each mitochondrion

While the hemibrain dataset comes with pre-saved files for the synapses and mitochondria of each neuron, this study required additional information to be saved for faster, later computations. For the mitochondria, this included the indices of which synapses are adjacent to each mitochondrion, the surface area of each mitochondrion, the cross-sectional area of each mitochondrion at its centroid, the circumference and cross-sectional area of the neuron at the centroid of the mitochondrion, the length of the mitochondrion along each principal component, the moment of inertia of the mitochondrion along each principal component, the symmetry of the mitochondrion along each principal component, the cross-sectional area and circumference of the mitochondrion using each principal component as the cross-sectional plane’s normal vector, the distance between the two farthest points on each mitochondrion, and the length of the mitochondrion along the instantaneous orientation of the neuron at the centroid of the mitochondrion (**Fig. 1c,d**; **Fig. 2a; Fig. S2**). Each of these values for each mitochondrion in the dendrite, connecting cable, and axon were stored in a CSV file for faster computation later.

The symmetry of the mitochondrion along each principal component is computed by first reflecting the mitochondrion about its centroid along the principal component, retaining all reflected and measured voxels in a new synthetic, symmetrized mitochondrion. The ratio of the symmetrized mitochondrion volume is then divided by the true mitochondrial volume to compute the symmetry metric, which is unity for perfect symmetry and increases for increasing asymmetry.

### Calculating the distance between mitochondria in the skeleton

To calculate the distance between objects in the neuron, we chose to compute the geodesic distance (the distance along the path of the neuron) instead of the Euclidean distance, sicne the Euclidean distance can differ substantially from the geodesic distance when it is greater than a few micrometers. For computing geodesic distance with synapses, we calculated the distance between the nearest nodes in the trimmed neuron skeleton and subtracted the distance from the synapse to their respective nearest node after projecting the synapse along the node’s orientation. However, mitochondria have a volume associated with them, so using the same methodology with their centroids would impose a minimum distance between two mitochondria, equal to the sum of half the length of each mitochondrion in the pair. Therefore, when calculating geodesic distances between mitochondria, we further subtract the overhang of the mitochondria on the skeleton node (**Fig. S1**). This is conceptually equivalent to subtracting half the length of the mitochondria, but it is accurate when mitochondria are curved. This allows the distance between mitochondria to be zero if they are directly adjacent to one other.

### Surface meshes

Surface meshes were computed for both mitochondria and the neurons to enhance visualization (**Fig. 1c**). This was done using the marching cubes algorithm ^84^.

### Coloring scheme

Colors for the LC neurons were chosen using the qualitative Brewer color scheme with seven groups ^85^. Given there were 21 LC neuron types, three different markers and line styles were used.

### Embedding mitochondria

All the mitochondria were embedded into two dimensions by their z-scored morphometrics using UMAP with default parameters (**Fig. 2b,d, Fig. S3**) ^43^. To visualize how mitochondria in each arbor are distributed in the embedding, we used kernel density estimation using the normal distribution approximation to compute the optimal bandwidth ^86^. The neuron types were sorted by hierarchical clustering to allow for easier visual comparison. The distance metric used for the covariance matrix was the average nearest neighbor distance in the full, z-scored morphometric space from the *i^th^* to *j^th^* neuron type.

### Random forest classifier

Random forest classifiers from the scikit-learn library ^87^ were used to classify LC mitochondria by which compartment they reside in (**Fig. 2c**), or which neuron type they came from (**Fig. 2f, Fig. S3**). In all cases, we used five-fold cross-validation when training the classifiers. When computing the Gini impurity, each mitochondrion is given a sample weight equivalent to the inverse number of mitochondria in the neuron type (for the neuron type classifier) or neuron compartment (for the compartment classifier) to prevent the classifier from being biased for neuron types with more mitochondria. Both classifiers used default parameters but were regularized by enforcing that each leaf in the decision trees contains at least 1% of the data. No hyperparameter search was done to find the optimal regularization strength.

Mitochondria were also classified into neuron types across the hemibrain (**Fig. S4**). For this model, a random forest classifier was trained to classify mitochondria into neuron types for all neuropils in the hemibrain that have at least 2 neuron types, each having at least 10 individual neurons. Each neuropil’s cell type classifier was evaluated on the mitochondria in each neuron type, for the mitochondria in the neuropil. To prevent overfitting, we used five-fold cross-validation and again enforced each leaf in the decision tree contains at least 1% of the data.

### Correlating differences in mitochondria morphology across neuron types for each cellular compartment

Within each of the three cellular compartments, the mean, z-scored morphometric features were calculated for each neuron type. The difference in mean morphometric features between a pair of neuron types for all three cellular compartments was compared by taking the cosine distance between the difference vectors for the dendrite and connecting cables (y-axis) and the axon and connecting cable (x-axis). These values are set as the y and x coordinates, respectively, for the scatter plots in **Fig. 2**.

Hypothetical distributions for various correlations in cell type specificity across cellular compartments were created *in silico* for illustrative purposes.

### Shuffling mitochondria

As a null hypothesis for many tests, mitochondria are shuffled to new locations in the neuron. When a mitochondrion is shuffled to a new location in the neuron, we place the mitochondrion in the same compartment (i.e. a mitochondrion in the axon is placed at a new, random location in the axon). The new location is a randomly chosen node in the trimmed, skeleton file. The coordinate of the new location becomes the center of mass of the shuffled mitochondrion. The morphometrics are preserved during shuffling, so all the morphometric statistics of the mitochondria pool in the compartments of the neurons are held constant.

### Positioning features

We computed features for both shuffled and measured mitochondria, characterizing the properties of the neuron and synapses near each mitochondrion (**Fig. 3a-c; Fig. S5**). These features can be grouped into two broad categories: histogram features and scalar/morphological features. The histogram features compute how many synapses or other mitochondria are in a bin at some distance away from the mitochondrion. The scalar/morphological features are a list of features that capture the morphology of the neuron near the mitochondrion.

For the histogram features, we find the geodesic distance from the mitochondrion to either the presynapses, postsynapses, or all the other mitochondria in the neuron. The 0 *um* bin simply counts the number of synapses with the mitochondrion, namely where the mitochondrion projection onto the skeleton overlaps the synapse projection onto the skeleton. The 0.3 – 0.6 *μm* bin feature counts the number of synapses or other mitochondria that are more than 0.3 *μm* away but less than 0.6 *μm* away. The other histogram features are computed similarly.

The scalar/morphological features were the “circumference retention”, “is on thicker daughter”, “leaf number”, “distance up section”, “branch order”, and “number of branches in”. The order of presentation of the scalar features in plots was chosen after fitting the logistic model from most to least predictive features (i.e. largest to smallest magnitude in logistic model coefficients).

- The “circumference retention” describes the narrowing of the neurite section a mitochondrion inhabits compared to its *mother section*. Mitochondria are always situated in *sections*, defined as the stretch between two branch points (where the neural arbor splits) or a branchpoint and a leaf node (i.e. a tip) of the neural arbor. Every section has a mother section, which is upstream, towards the direction of the soma; sections downstream from the mother are its daughters. The circumference of a section is the average circumference of all the SWC file nodes whose cross-sectional plane does not intersect any mitochondrial tissue. Points with mitochondria are ignored when computing circumference, because there are often bulges where mitochondria are located. Finally, circumference retention is defined as the ratio of the circumference of the section with the mitochondrion divided by the circumference of its mother section.
- The “is on thicker daughter” feature is a Boolean feature that compares the circumference of the section with the mitochondrion and the circumference of its sister section, defined as the other daughter section of its mother section.
- The “leaf number” is the number of leaf nodes that are downstream (away from the soma) from the mitochondrion.
- The “distance up section” is computed by first calculating the total geodesic length of the section the mitochondrion is in. Next, the geodesic distance from the mitochondrion to the upstream branch point (towards the soma) is divided by the total geodesic length of the section to get the distance up the section.
- The “branch order” is the number of branch points between the mitochondrion and the base of the arbor the mitochondrion is in.
- The “number of branches in” is the integer number of branch points the mitochondria is in. For instance, if the mitochondrion is very small and in the middle of a segment, it will likely not be in any branch points. However, if the mitochondrion is very long, it may travel through multiple branches. Additionally, some mitochondria are positioned right at a branch point, so those mitochondria are in one branch.

### Mitochondria positioning logistic model training

Generalized linear models were trained (using the Python statsmodels library ^88^) to estimate the probability a mitochondrion is in its measured position in the neuron (**Fig. 3d,i,j; Fig. S5**).

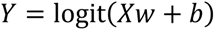

Where *Y* is the probability that the mitochondrion is present, *w* is the learned weights, *X* is the data matrix of positioning features for the measured and shuffled mitochondria, and *b* is the learned bias. The logit link function was used with a cross-entropy loss function. To prevent overfitting, we used 5-fold cross-validation and lasso regression with varying regularization strengths until the maximum test AUC was achieved. Then, the model was retrained without any regularization, using only features with non-zero coefficients from the regularized model.

### Jittering mitochondria

Mitochondria were jittered by moving their center of masses either up or down the neuron based on a Gaussian distribution with varying standard deviations (**Fig. 3e-g**). This is computationally analogous to moving the mitochondria along the cytoskeleton in the anterograde or retrograde direction, respectively. When the mitochondria’s center of mass is moved along the skeleton, it may move past the base of the arbor or a leaf node. In this case, the mitochondria are reflected in the opposite direction. Whenever a jittered mitochondrion crosses a branch point, we set it to have an equal probability of moving into the two available branches. Mitochondria have an equal probability to initially move up or down the arbor. Once all the mitochondria are jittered, the mean absolute displacement was computed from the original mitochondria locations to their respective new locations after the jitter process. The reflective boundary conditions enforce an upper bound on the mean absolute displacement of the mitochondria.

### Dimensionality reduction for positioning rules

Logistic model weights were projected into the first two principal components of the weights across the different models to visualize differences in the rules between neuron types and axons/dendrites (**Fig. 3j**). For each model, confidence interval ellipses were projected into the same principal component plane from the parameter covariance matrix for each model, *C*_*i*_, computed by the statsmodel library ^88^.

To judge differences between pairs of rules, we computed the Euclidian distance between logistic model weights divided by the standard deviation of the parameter fit in the direction of the difference. To find this, we computed the number of standard deviations between the *i^th^* and *j^th^* models as 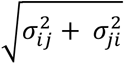, where *σ_ij_* is the confidence interval on the parameters of the *i*th model projected into the direction of the vector connecting the *i*th and *j*th model parameters, that is, *σ*_ij_ = 〈*θ*_*_ − *θ*_i_|*C*|*θ*_j_ – *θ*_i_〉, where *θ_i_* is the parameter set for the *i*th model and *C_i_* is the parameter covariance matrix for the *i*th model.

### Connectivity and functional metrics

Four metrics were used to quantify the connectivity and functional properties of each LC neuron type: Lobula innervation distance, broad connectivity distance, glomeruli distance, and calcium response distance (**Fig. 4e-i**). The lobula is a neuropil in the optic lobe, where each LC dendritic arbors innervate. LC neurons innervate different layers of the lobula, which has consequences for which upstream neurons will synapse onto each LC neuron (Nern, Loesche et al. 2024). The depth axis of the lobula penetrates through these layers and was set to be the third principal component of the lobula synapses of the LT1 cell type (Tanaka and Clark 2022). The lobula innervation distance between neuron types was computed as the cosine distance of the histogram of dendritic synapses along the depth axis of the lobula.

The broad connectivity distance compared the downstream connectivity of different neuron types. Since each LC neuron type has unique postsynaptic targets, the downstream neuron types were grouped by their class. The class was defined as the list of letters at the beginning of the neuron type name. For instance, PVLP028 and PVLP115 would be grouped as PVLP. The broad connectivity distance between neuron types was therefore the cosine distance between the average downstream connectivity vector (number of connections onto each downstream neuron class) for each neuron type.

The glomeruli distance compares the location of the axons of each neuron type. Glomeruli were defined as all the axonal synapses of a given neuron type. The glomeruli distance was the Euclidean distance between the centroids of each pair of glomeruli.

The calcium response distance builds on prior work ^39^, which recorded the calcium response of several LC neuron types to a suite of visual stimuli. In that study, the responses to all visual stimuli for each recording were normalized to be between 0 and 1 (by subtracting the minimum and then dividing by the resulting maximum). We followed that work to compute the functional distances in the plane of the first two principal components: after projecting the normalized recordings into their first two principal components, the calcium response distance was defined as the Euclidean distance of the centroids for a pair of neuron types in the two-dimensional space.

Each of these metrics was correlated with the positioning rule distance (**Fig. 4j,k; Fig. S6**). The positioning rule distance is defined as the Euclidian distance between logistic model weights, *W*, for the two neurons.

### Enrichment of presynapses with mitochondria

Synapses are referred to as being “with mitochondria” if the cross-section of the neuron at the coordinates of the synapse has any mitochondrial tissue. The “mitochondrial coverage” is defined as the fraction of presynapses with mitochondria, *f*. Coverage can be computed for all synapses or subsets of synapses based on different criteria, *f*_*i*_, typically by conditioning on the postsynaptic partner. We computed enrichment, *E*_*i*_, of coverage by comparing one set of presynapses (for instance, with a specific postsynaptic partner) to all the remaining presynapses (for instance, with all other postsynaptic partners). In particular, though we compared distributions in various ways, we computed the enrichment as the ratio of *f*_*i*_ to *f* computed from the remaining synapses.

When computing enrichment by postsynaptic neuron type for the LC neurons, we included in our analysis all postsynaptic neuron types that have at least 5 connections for at least half of the LC neuron type’s neurons (**Fig. 5c; Fig. S7**). To compute the enrichment, we compared mitochondrial coverage for presynapses with one postsynaptic partner, *i*, to all other presynapses.

Variability in the enrichment for a neuron type, *k*, was computed by calculating the root mean square enrichment of the measured dataset, then subtracting an equivalent measurement for version of the dataset where the connectivity of the presynapses was shuffled (**Fig. 5d**). The variability was thus 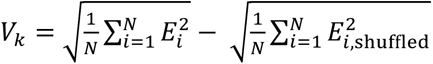, where *E*_*i*_ is computed for presynapses in neuron *k* and *N* is the number of postsynaptic partners.

For the Kenyon cell (KC) neuron types, all KCs with at least 10 individual cells were used. To analyze how mitochondrial coverage of synapses differs between mushroom body compartments, the fraction of presynapses with mitochondria in each mushroom body compartment (test condition) was compared to the fraction of presynapses with mitochondria in all other compartments (**Fig. 5f,h,j; Fig. S8**). Only compartments that have at least 5 presynapses for at least half of the KC neuron type’s neurons are shown. To analyze how the mitochondrial linear density varies across mushroom body compartments, we computed the linear density as the total length of all mitochondria in the mushroom body compartment divided by the total neurite length in the mushroom body compartment (**Fig. 5g,h; Fig. S8**). To obtain the enrichment in each compartment, we compared the linear density in each compartment to the linear density in the other compartments. When computing enrichment of coverage based on postsynaptic targets, we included postsynaptic targets that received at least 5 connections from at least half of the KC neurons in the KC neuron type and comprise at least 5% of the total heterotypic connections from the KC neuron type (**Fig. 5i**).

In all cases, to find enrichment, we computed differences in a pairwise manner, within each cell. Error bars for the enrichment calculations are found by resampling the dataset with replacement and then finding the standard deviation over the distribution of resamplings.

---

**Figure S1.**
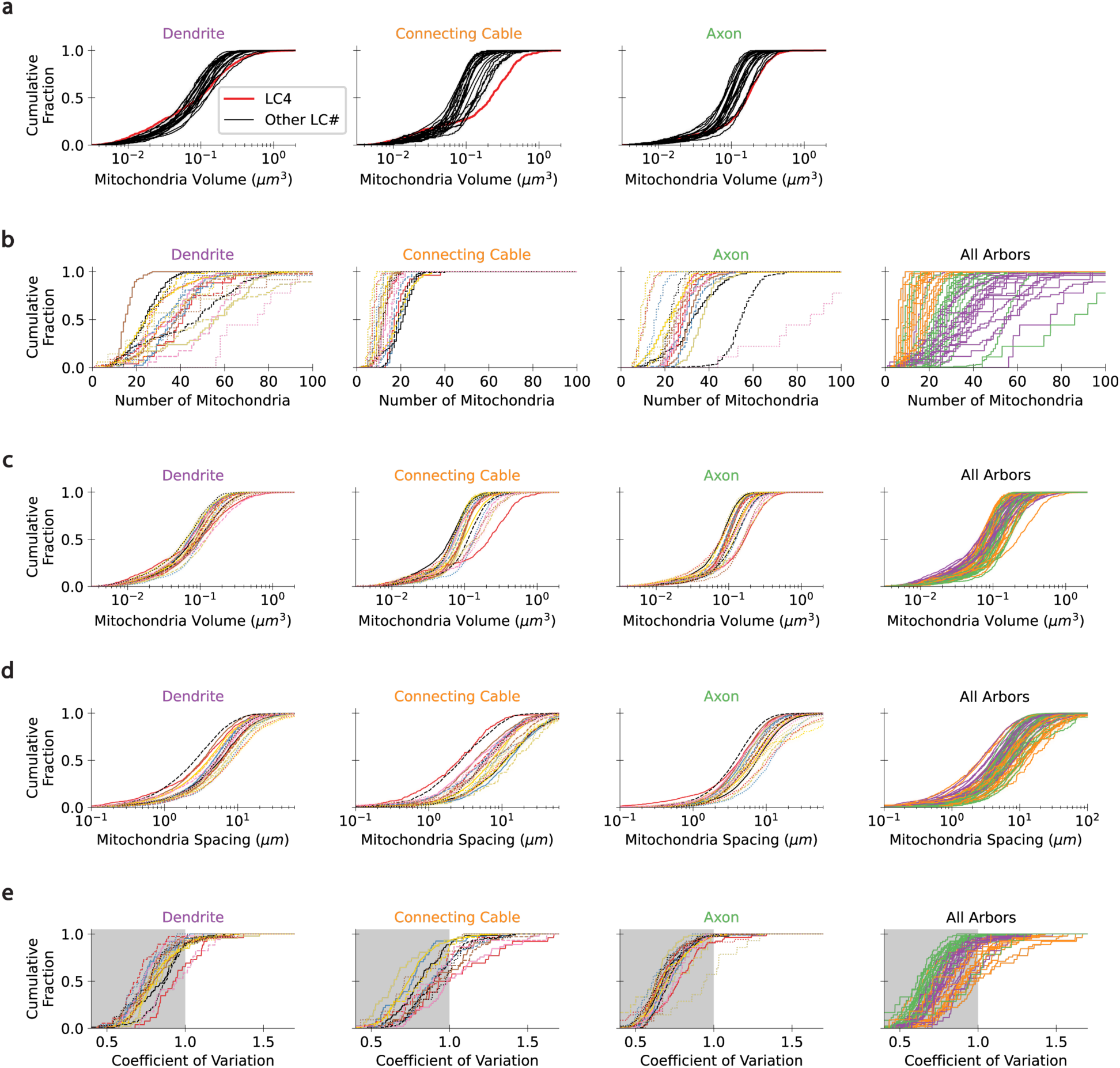
Distributions of mitochondria statistics across LC neuron types and segments. (a) Cumulative distribution functions of mitochondria volumes for all LC neuron types in the dendrite (left), connecting cable (middle), and axon (right). The CDF for LC4’s mitochondria in each cellular segment is labeled red. (b,c,d,e) Cumulative distribution functions for the number of mitochondria (b), mitochondria volume (c), mitochondria spacing (see **Methods**) (d), and coefficient of variation of mitochondria spacing (e) for each LC neuron type in each segment. All segments are plotted on the right with the segment color scheme.

**Fig S2:**
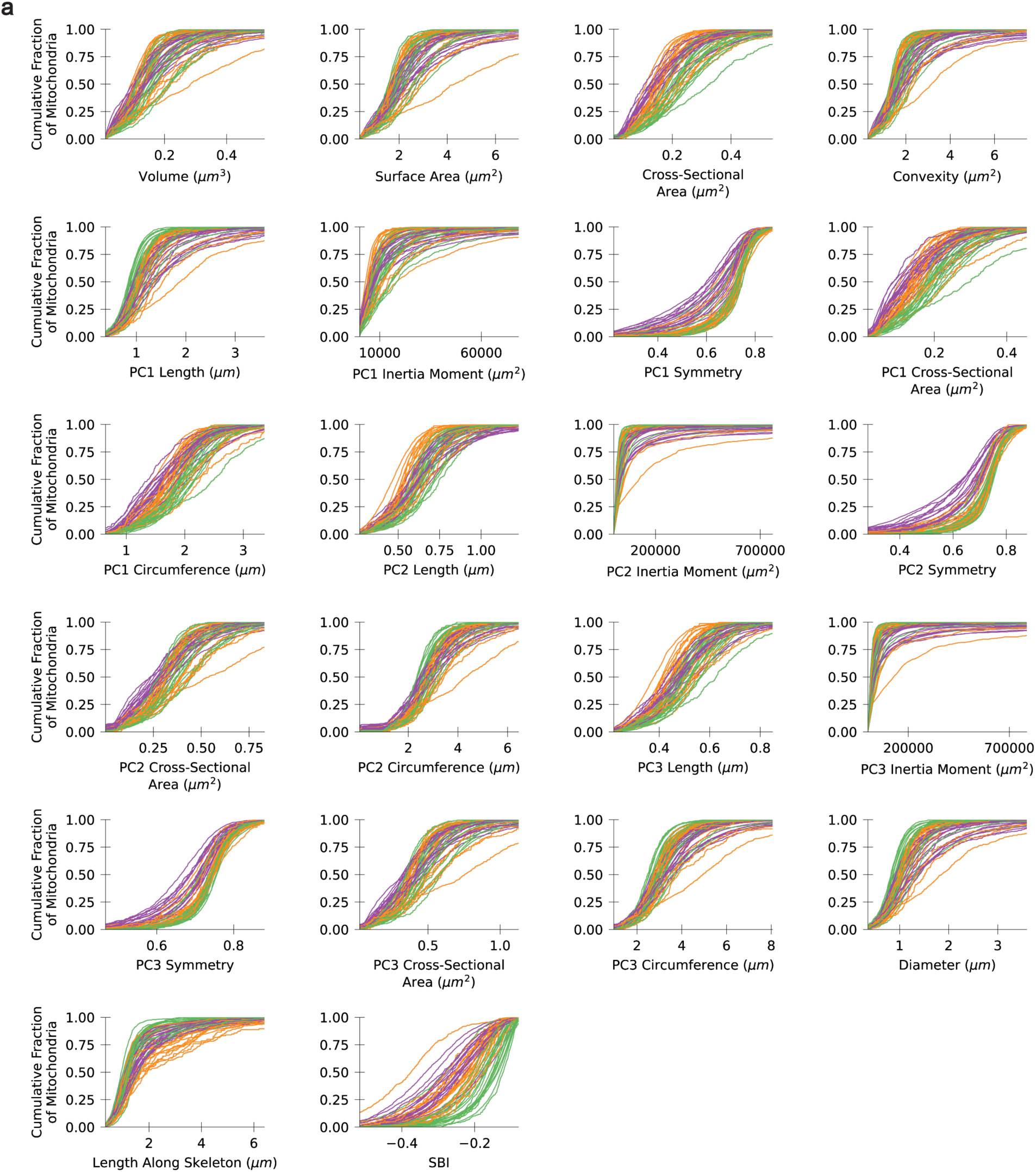
Mitochondrial morphometric distributions. (a) Cumulative distributions for all the mitochondrial morphometric features (not z-scored) in each LC neuron type and each segment. Axons are green, connecting cables are orange, and dendrites are purple.

**Figure S3.**
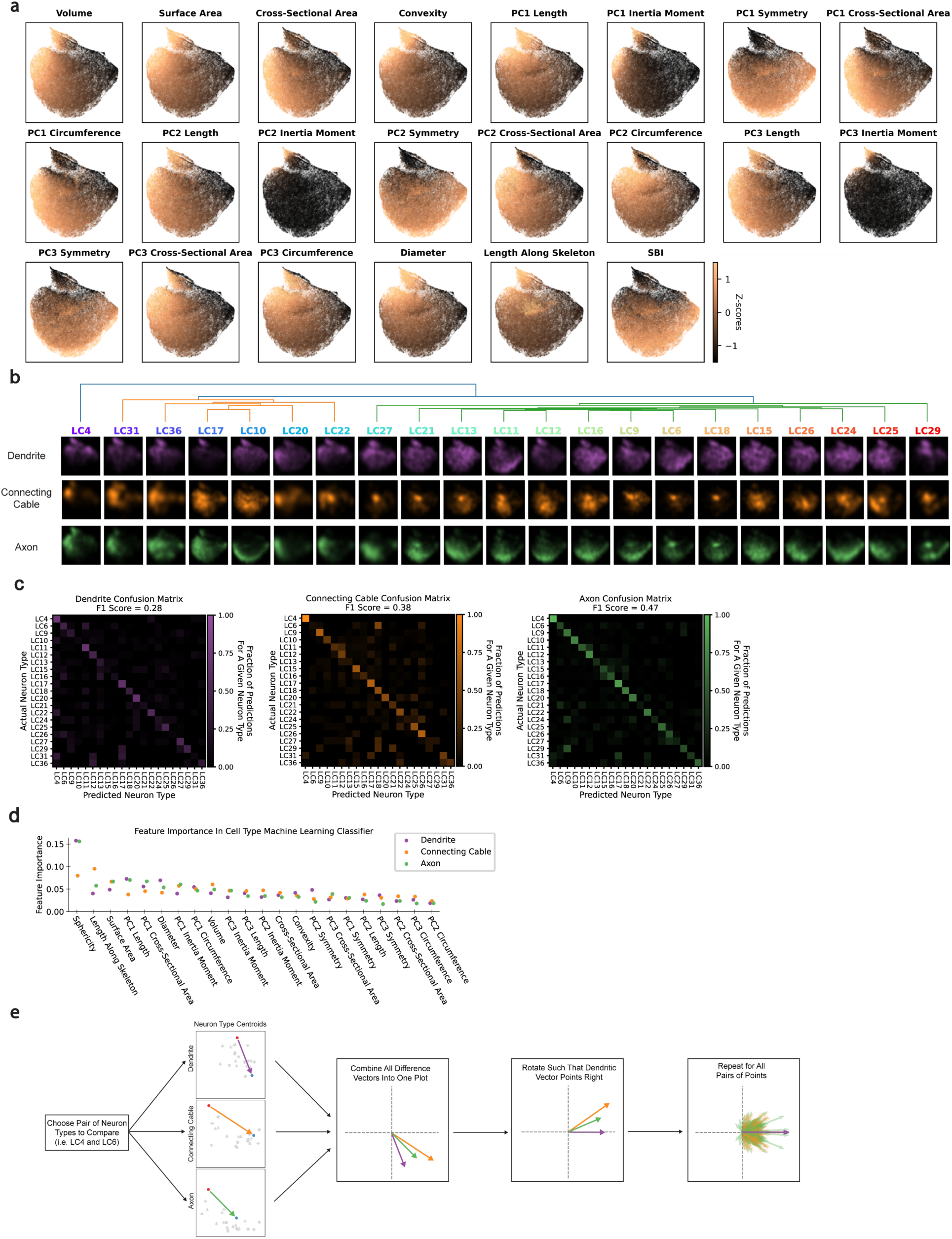
Morphometric features of mitochondria are separated by UMAP embedding. (a) UMAP embedding of the z-scored mitochondria morphometrics, as in Figure 2b, but points are colored by input feature values. Each plot colors the points according to the z-score of the point each feature. See methods for how morphometric features are calculated. (b) Kernel density estimates of all LC neuron type mitochondria embedding in each segment. The dendrogram shows a hierarchical clustering of all LC neurons types by their mitochondria morphometrics. (c) Confusion matrices when using trained random forest models to predict the cell type from using only the dendritic (left), connecting cable (left middle), or axonal (right middle) mitochondria. (d) Feature importance of each morphometric dimension used in the random forest classifier in Fig. 2f. (e) Cartoon illustrating how differences in mitochondria morphology between cell types were qualitatively compared across cellular compartments.

**Figure S4.**
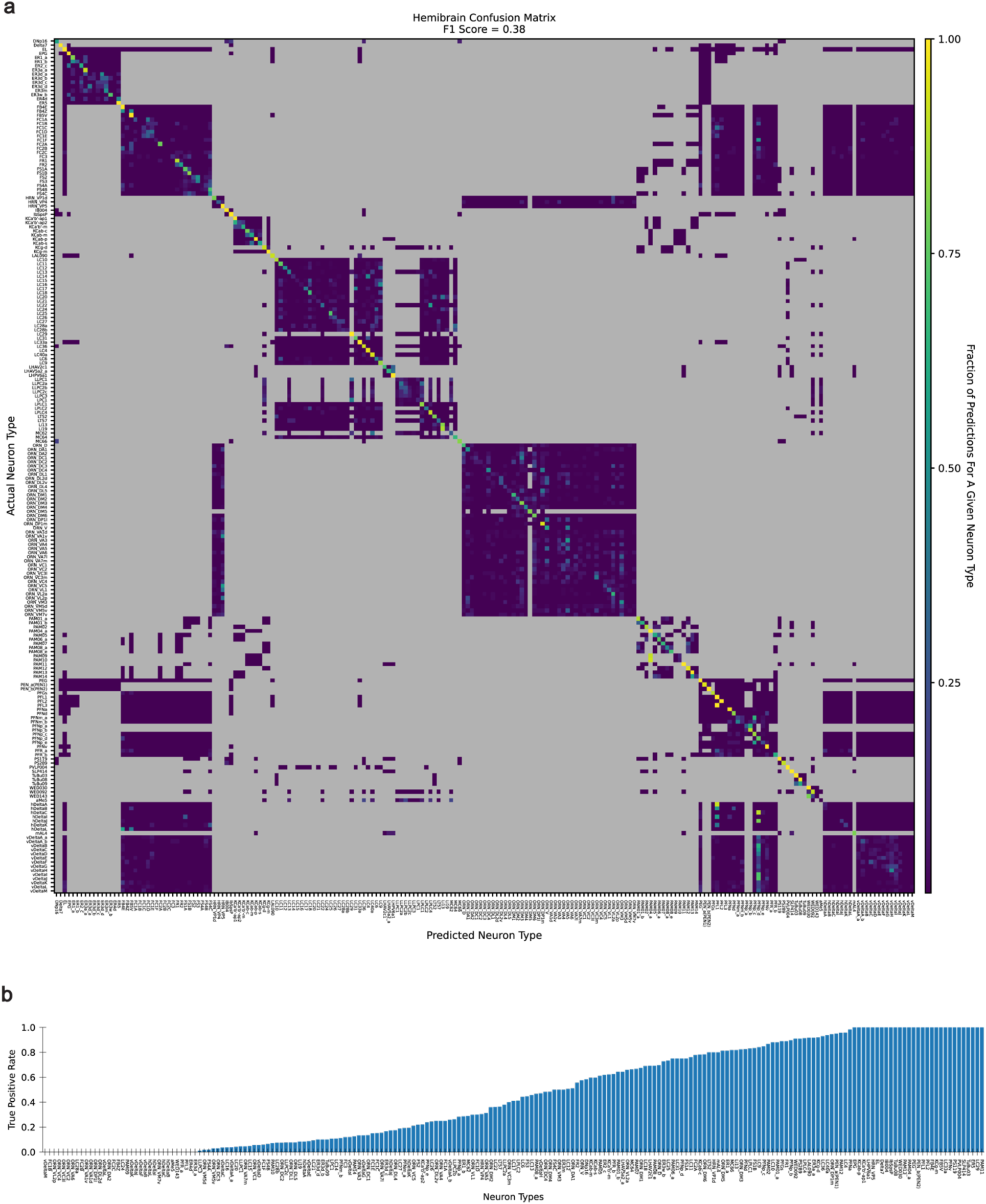
Neuron type classifier across hemibrain neurons. (a) Confusion matrix for a whole hemibrain neuron type random forest classifier, which uses the morphometric features of each neuron’s mitochondria in each neuropil. All neuron types with at least 10 neurons in the hemibrain were included. In the confusion matrix, intersections between neurons that do not innervate a similar neuropil are colored grey, since the classifier is trained and tested for each neuropil individually. Thus, these gray patches correspond to predicted fraction of 0, but simply because the neurons cannot be confused with one another because they do not share neuropil. (b) True positive rates for the neurons in (a).

**Figure S5.**
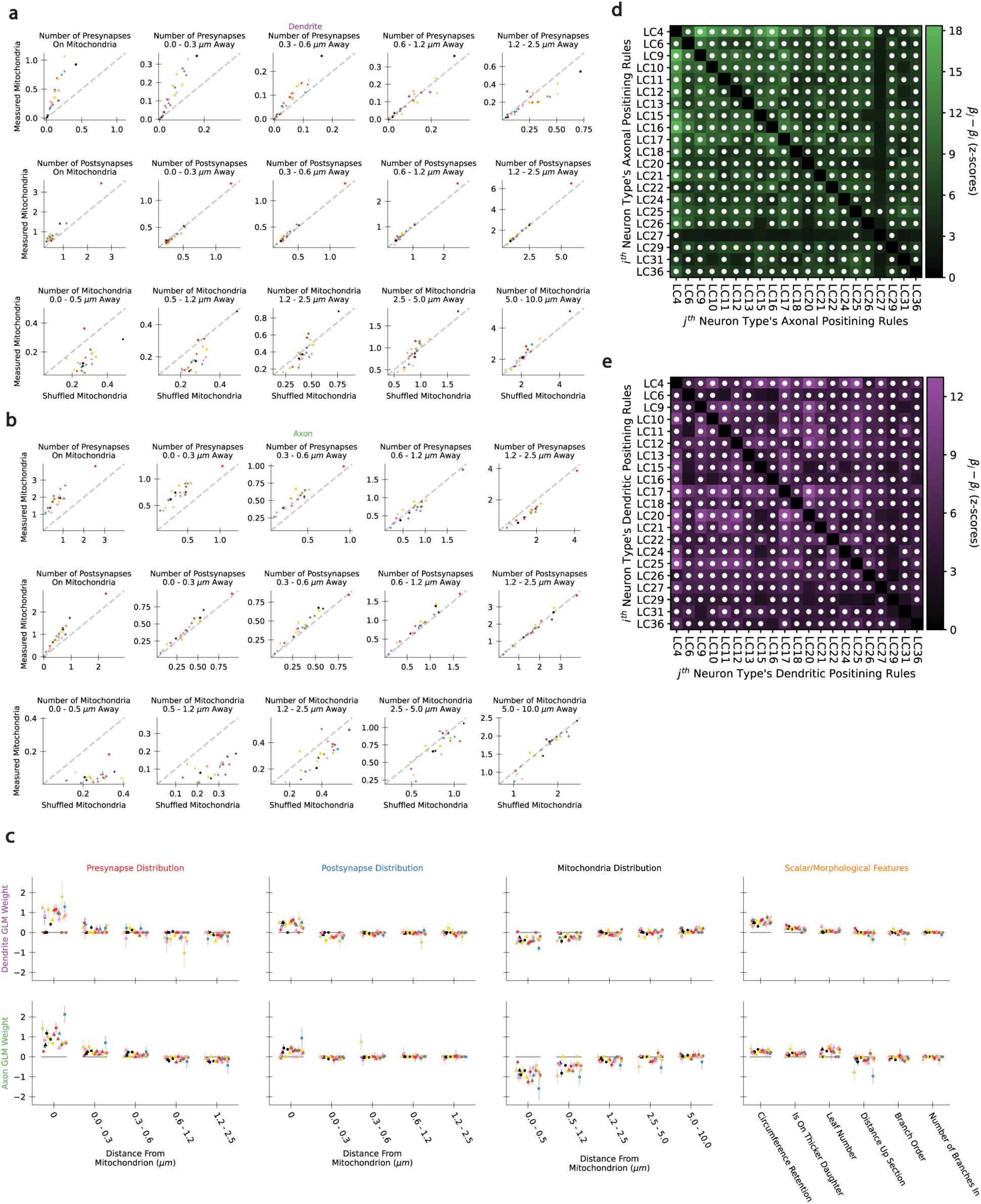
Measured mitochondria positioning features differ from features with randomly positioned mitochondria. (a,b) For each feature used in the logistic models in Figure 3, the mean value of the feature in the dataset is plotted against the mean value of the feature for a dataset that shuffled mitochondria locations. The is plotted for the dendrite (a) and axon (b) of every LC neuron type. The grey dashed line is the unity line. Points above the unity line indicate a positive correlation of mitochondrial placement with that feature, while points below the line indicate a negative correlation. (c) Plot of model weights for the logistic positioning model for all LC neuron types, showing error bars for the 95% confidence interval. See methods for descriptions of the features. (d,e) Matrix for all pairwise differences in the logistic model weights for all LC neuron types in their dendritic (d) and axonal (e) models, units of z-scores (see **Methods**). Matrix elements with statistically significant values after Bonferroni correction are labeled with white dots.

**Figure S6.**
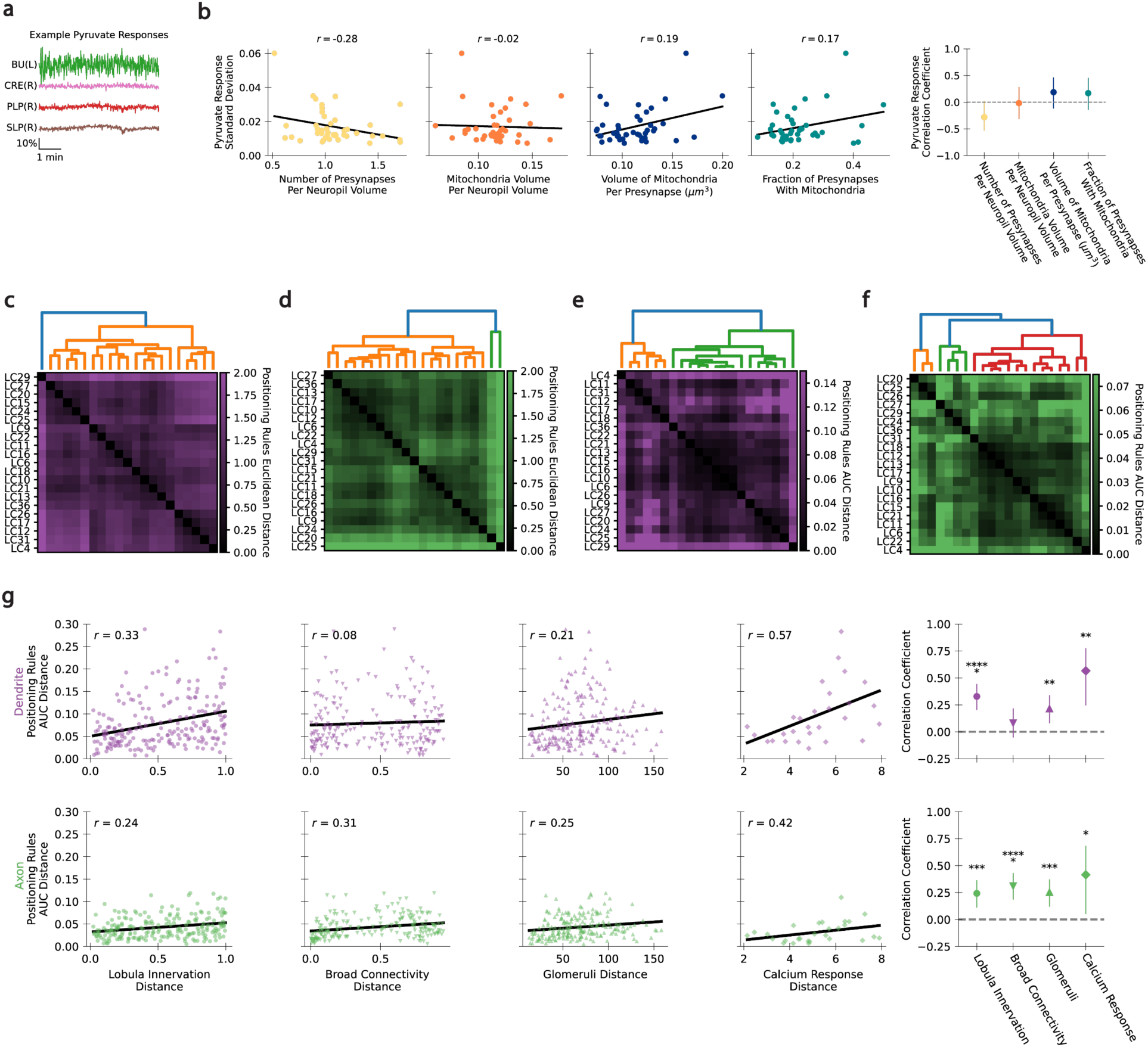
Mitochondrial positioning rules correlate with function and connectivity. (a) Example pyruvate indicator traces for four central brain neuropils (data from ^50,51^). (b) Scatter plots of the standard deviation in the pyruvate indicator responses for all recorded central brain neuropils against the number of presynapses per neuropil volume (left), total volume of mitochondria per volume of neuropil (left middle), total volume of mitochondria per presynapse (right middle), and fraction of presynapses with mitochondria (right). Best fit lines are in black. The Spearman correlation coefficients with 95% confidence intervals are at the far right. (c,d) Distance matrix for the all pairwise positioning rules distances between all LC neuron types for the (c) axonal and (d) dendritic arbors. Distance is computed as the square root of the sum of square differences in the logistic model weights. (e,f) Distance matrix for the all pairwise positioning rules distances between all LC neuron types for the (e) axonal and (f) dendritic arbors. Distance is computed by summing of the AUCs when a model was trained on one neuron type and tested on the other neuron type (and vice versa), then subtracting the sum of the AUCs for each neuron in the pair when the logistic model is trained and tested on the same neuron type. (g) Scatter plot of positioning rules AUC distance with each functional and connectivity distance for all pairs of LC neuron types, for dendrites (top) and axons (bottom). Black line represents the best fit line. The right panels are the Spearman correlation coefficients with 95% confidence intervals. P-value significance is represented by asterisks (* < 0.05, ** < 0.01, *** < 0.001, with additional asterisks for additional factors of 10).

**Figure S7.**
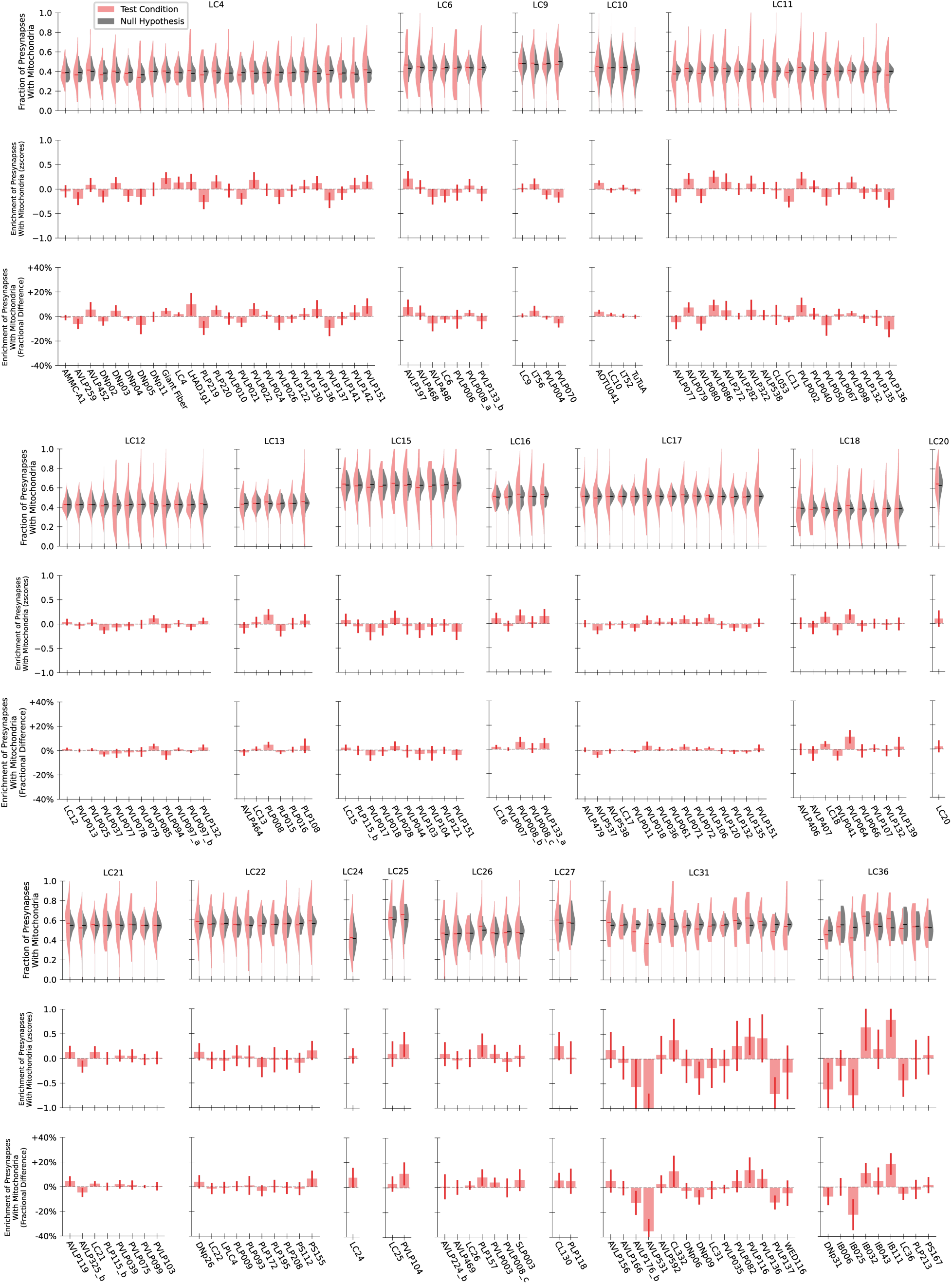
Presynaptic mitochondrial coverage in LC neurons shows minimal enrichment or depletion for different postsynaptic neuron types. For every LC neuron type, the kernel density estimates (top row) of the presynaptic mitochondrial coverage for a particular postsynaptic neuron (test condition) is compared to all other presynapses (null hypothesis). The distributions are over all cells of that cell type. The enrichment of presynapses with mitochondria for each postsynaptic connection is also shown in units of z-scores (middle row) and fractional differences (bottom row).

**Figure S8.**
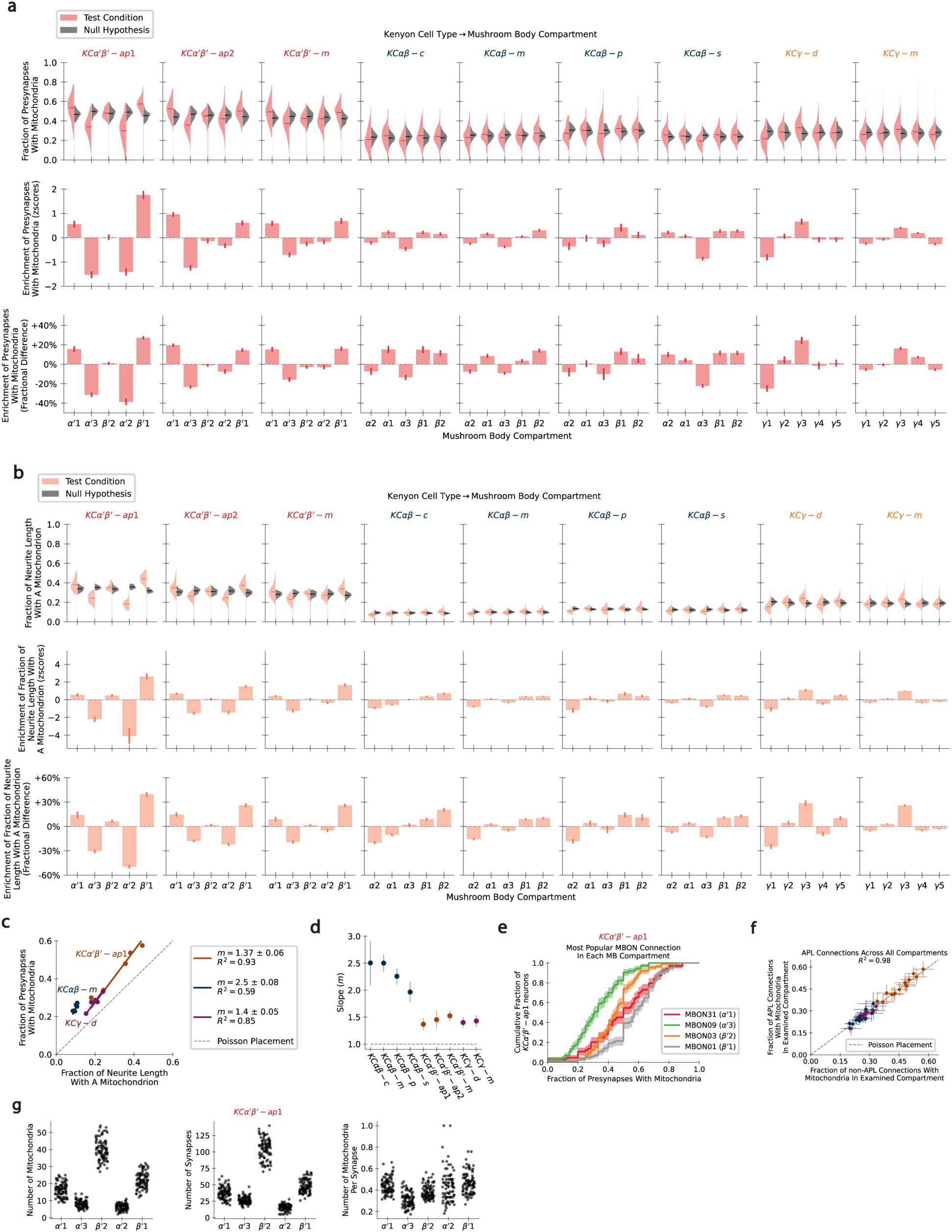
Mitochondrial coverage in Kenyon cells depends on the functional compartment. (a) For every KC neuron type, the kernel density estimate (top row) of the fraction of presynapses with mitochondria for presynapses in a particular mushroom body compartment (test condition) are compared to all other presynapses (null hypothesis). The enrichment of presynapses with mitochondria in each mushroom body compartment is shown in units of z-scores (middle row) and fractional difference (bottom row). (b) For every KC neuron type, the kernel density estimates and enrichments of the fraction of neurite length with a mitochondrion (i.e., linear mitochondrial density) are shown. This is computed as in (a). (c) Scatter plot of the fraction of presynapses on mitochondria against the fraction of neurite length with a mitochondrion (mitochondrial linear density) for each mushroom body compartment, for three representative KC neuron types. Points for each KC type are fit to a line with the slope (m) and coefficient of determination (*R*^2^). (d) Plot of slopes in (c) with 95% confidence intervals for all Kenyon cell types. Each neuron type is colored by its Kenyon cell class as in Fig. 5e. (e) Cumulative distribution function of *KCα*′*β*′ − *ap*1 presynapses with mitochondria for presynapses connecting onto the MBON with the most connections in each compartment. (f) For every mushroom body compartment that each of the nine KCs innervate, the fraction of APL presynapses with mitochondria is plotted against the fraction of presynapses with mitochondria in that compartment, excluding APL presynapses. The grey dashed line represents the expected result if mitochondrial coverage depends on the mushroom body compartment. (g) Plots of the number of mitochondria (left), number of synapses (middle), and number of mitochondria per synapse (right) in each *KCα*′*β*′ − *ap*1 in each mushroom body compartment it innervates.

**Figure S9.**
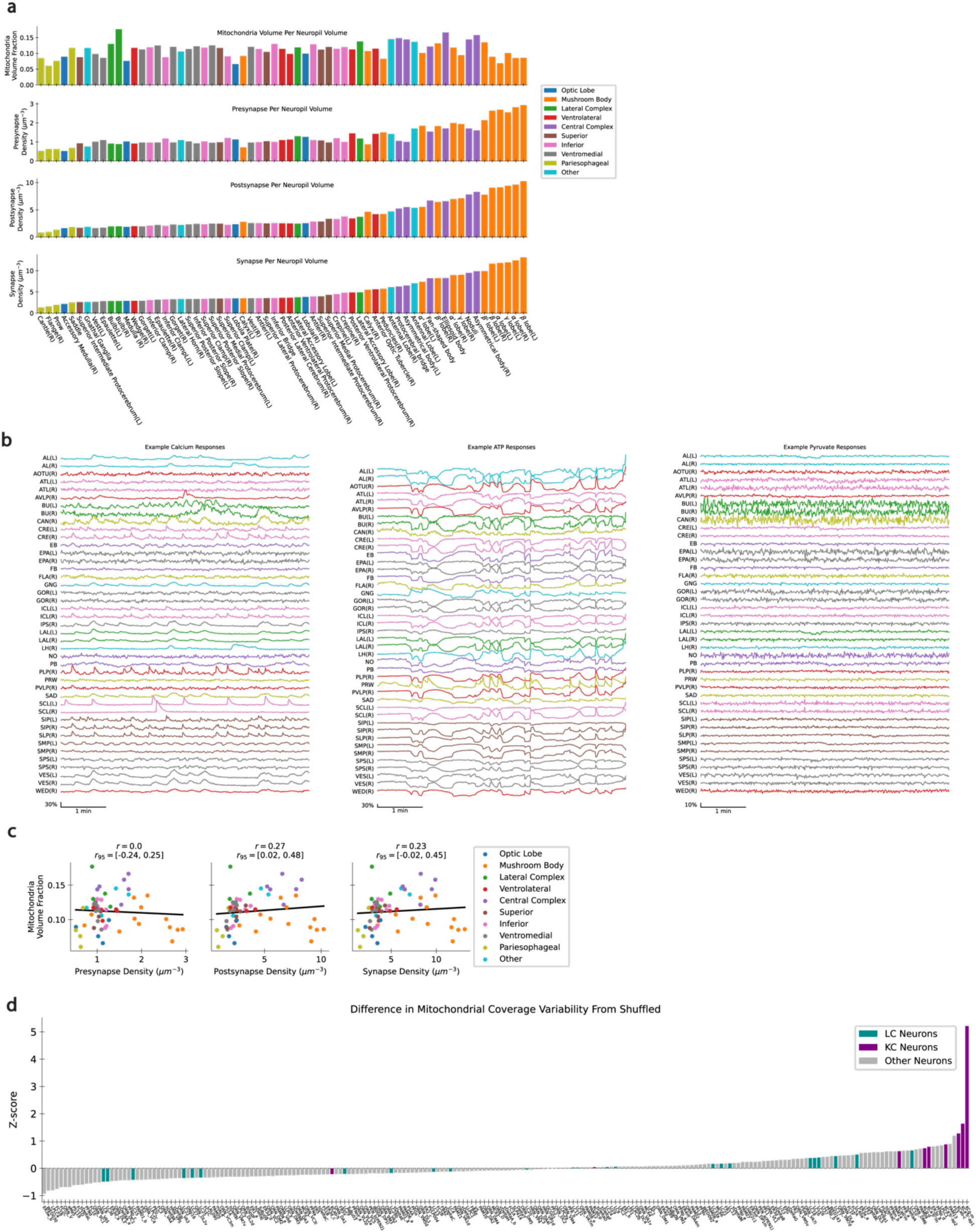
Mitochondria statistics across the hemibrain. (a) Plots of the mitochondria volume per neuropil volume (top), number of presynapses per neuropil volume (top middle), number of postsynapses per neuropil volume (bottom middle), and total number of synapses per neuropil volume (bottom). Neuropils are colored by their class and are sorted in increasing order of total number of synapses per neuropil volume. (b) Example calcium, ATP, and pyruvate indicator traces from every central brain neuropil (data from ^50,51^). (c) Correlation between the mitochondria volume fraction with the presynapse density (left), postsynapse density (middle), and total synapse density (right). Each point is a distinct neuropil and is colored by its respective class, as in (a). The Spearman correlation coefficient and 95% confidence interval are above each plot. (d) Difference in the mitochondrial coverage variability from a shuffled null hypothesis as also plotted in Figure 5d, but here in units of z-score, showing how distinguishable each neuron type’s coverage variability is from the shuffled null hypothesis. All neuron types in the hemibrain with at least 10 individual neurons are shown. Neuron types are sorted in order of increasing z-score. Note that the KCs are highly distinguishable from random in part because there are many neurons in each type.

**Figure S10.**
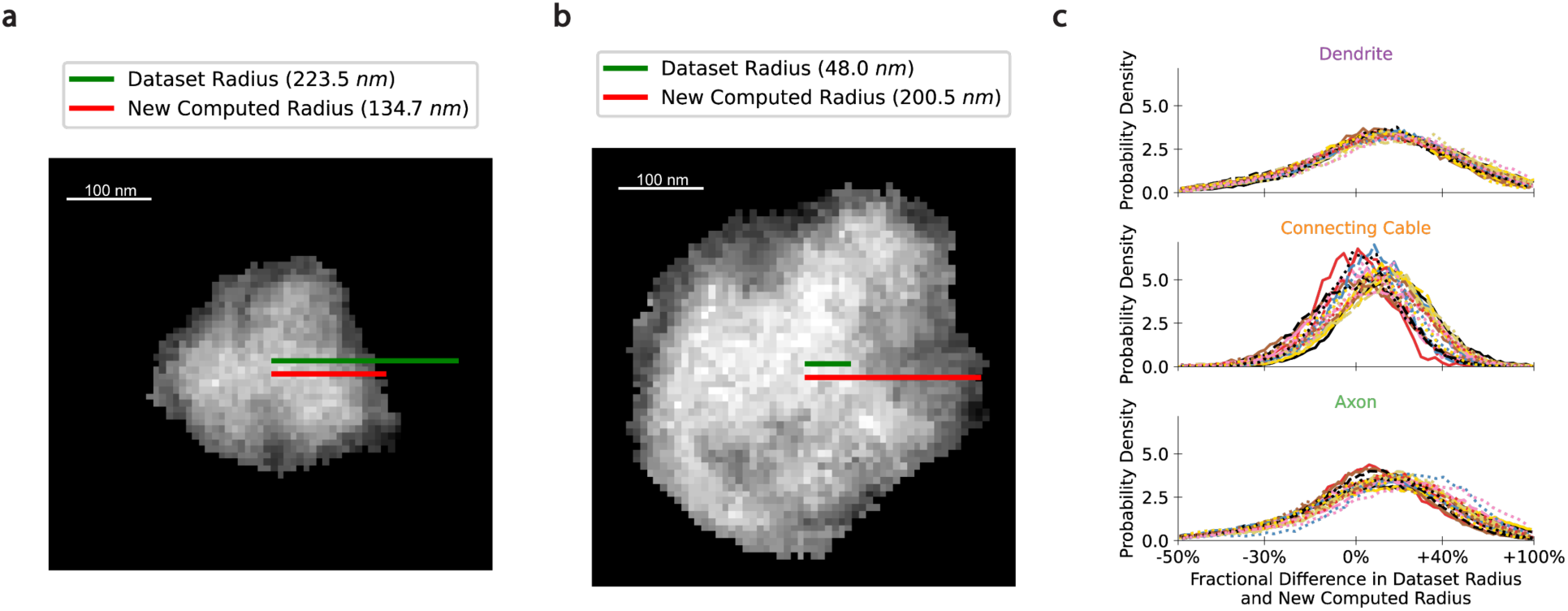
Newly computed neurite radii are up to 50% different from those listed in the dataset. (a) Example neuron cross-section of the axon of an LC12 neuron (bodyId 1343378332). The red line from the center of mass represents the newly computed radius (see **Methods**), and the green lines represent the radius listed in the dataset. (b) Example neuron cross-section of the axon of an LC9 neuron (bodyId 1229507904). Red and green lines as above. (c) Distribution of the fractional difference of the dataset radii from the newly computed radii for all 21 LC neuron types in each of the three compartments. The radii in the dataset can vary from the new computed radii by up to 50%.

